## Supplementary Information for "Transertion and cell geometry organize the *Escherichia coli* nucleoid during rapid growth"

Supplementary Methods

Supplementary Notes 1-2

Supplementary Figures 1-23

Supplementary Tables 1-4

Supplementary Videos 1-3 (Stills)

### Supplementary Methods

#### Generation of chromosomal fluorescent protein fusions

*E. coli* strain **CS1** (*E. coli* MG1655 chromosomally expressing MreB$^{sw}$-sfGFP and HupA-mCherry) was constructed using λ-red recombineering. It carries a fluorescent protein fusion of the cytoskeletal protein MreB and msfGFP (parental strain NO34 [1]) and a fluorescent protein fusion of the nucleoid-associated protein HU-alpha and mScarlet-I. For recombineering, plasmid pKD46 was transformed into electro-competent NO34. The DNA fragment for C-terminal insertion of mScarlet-I including a flexible linker (GSAGSAAGSGEF) and a chloramphenicol resistance cassette was generated as follows: Fragment 1 contains a 147 bp overlap to the C-terminal sequence of *hupA* and the linker. It was amplified from the NO34 genome using primers CS_FFM_002 and CS_FFM_003. Fragment 2 (linker-mScarlet-I) was amplified from Addgene plasmid #85044 (pmScarlet-I_C1) [2] using primers CS_FFM_005 and CS_FFM_006. Fragment 3 including the Chloramphenicol resistance cassette and a 59 bp overlap to the downstream region of *hupA* (mScarlet-I-FRT-CAT-FRT-hupA_ds) was amplified from Addgene plasmid #101148 (pmMaple3-CAM) [3] using primers CS_FFM_017 and CS_FFM_018. Fragments 1 and 2 were fused in a 2-step PCR using amplification primers CS_FFM_002 and CS_FFM_006, and the resulting fragment was fused similarly with fragment 3 using amplification primers CS_FFM_002 and CS_FFM_019. The resulting DNA fragment (1857 bp) was electroporated into NO34 carrying pKD46. Clones carrying the insertion were selected on plates containing Chloramphenicol and clones were verified by sequencing and fluorescence microscopy.

*E. coli* strain **CS2** (*E. coli* MG1655 chromosomally expressing MreB$^{sw}$-sfGFP and H-NS-mCherry) was constructed similar to CS1. It carries a fluorescent protein fusion of the cytoskeletal protein MreB and msfGFP (parental strain NO34 [1]) and a fluorescent protein fusion of the nucleoid-associated protein H-NS and mScarlet-I. The DNA fragment for C-terminal insertion of mScarlet-I including a flexible linker (GSAGSAAGSGEF) and a chloramphenicol resistance cassette was generated as follows: Fragment 1 contains a 155 bp overlap to the C-terminal sequence of *hns* and the linker. It was amplified from the NO34 genome using primers CS_FFM_012 and CS_FFM_013. Fragment 2 is identical to the fragment used for construction of CS1. Fragment 3 including the Chloramphenicol resistance cassette and a 54 bp overlap to the downstream region of *hns* (mScarlet-I-FRT-CAT-FRT-hns_ds) was amplified from Addgene plasmid #101148 (pmMaple3-CAM) using primers CS_FFM_017 and CS_FFM_020. Fragments 1 and 2 were fused in a 2-step PCR using amplification primers CS_FFM_012 and CS_FFM_006, and the resulting fragment was fused similarly with fragment 3 using amplification primers CS_FFM_012 and CS_FFM_021. The resulting DNA fragment (1860 bp) was electroporated into NO34 carrying pKD46. Clones carrying the insertion were selected on plates containing Chloramphenicol and clones were verified by sequencing and fluorescence microscopy.

*Xenorhabdus doucetiae* strain **CS_Xd1** was constructed using a pDS132-based suicide plasmid. First, a pCK-MreB-sfGFP plasmid (pDS132 with chloramphenicol resistance cassette) was constructed. The plasmid contains the *X. doucetiae* MreB gene with an msfGFP inserted into an internal loop between amino acids T228 and D229. For the construction of the plasmid, Fragment 1 (left homology region of MreB) was amplified using primers CS_MPI_003 and CS_MPI_004. Fragment 2 is the msfGFP sequence (*E. coli* codon-optimized) flanked by two linker regions ('SGSS' on the left and 'SGAPG' on the right) and was amplified from genomic DNA of the *E. coli* strain NO34 using primers CS_MPI_018 and CS_MPI_019. Fragment 3 contains the remaining sequence of the MreBCD operon and was amplified with primers CS_MPI_005 and CS_MPI_006. As backbone, plasmid pCK-CipA (suicide vector pDS132 derivative with Chlorampenicol resistance cassette) was digested with PstI and BglII (NEB). Fragments and backbone were assembled into one plasmid using Gibson Assembly (NEB HiFi DNA assembly kit) and electroporated into competent cells of the *E. coli* ST17 λ-pir conjugation strain. Clones were selected on chloramphenicol-containing agar plates and verified by colony PCR using primers VpDS132_fw and VpDS132_rv. The plasmid was transferred into *X. doucetiae* WT cells via conjugation. An *X. doucetiae* clone with chromosomally integrated pDS132 plasmid was grown overnight without antibiotics and 5 µl were streaked on LB agar plates containing 6% sucrose for the second recombination. Clones were screened by fluorescence and insertion of GFP at the proper site was confirmed by sequencing. PCR templates for sequencing were amplified using primers CS_MPI_020 and CS_MPI_021.

#### Supplementary Notes

##### Supplementary Note 1: The effect of 2D projection in CLSM and SMLM images

Projection effects are omnipresent in 2D imaging [4], with the only notable exception of TIRF imaging. Fluorescence signal detected within a certain 3D volume is projected onto a 2D plane on the sensor (here the camera chip), providing a distorted view of the target's true localization and spatial distribution. The sphero-cylindrical shape of gram-negative bacteria is hereby particularly problematic, as signal above or below the centre plane is projected into the central part of the 2D cross-section. In SMLM, this effect can be reduced by filtering single-molecule localizations by their width, as PSF width increases with axial distance from the focal plane [5]. The resulting projection depth, however, still comprises several hundreds of nanometres. Practically, this results in nucleoids that appear to span the cell cytosol. While this is might be the case occasionally, most cytosol-crossings of nucleoids during fast growth are likely 2D projection artefacts (see **Figure 1**). In widened cells or bacteria with larger diameters, parts of the membrane-attached nucleoid reside outside of the projection volume. As the projection volume is defined by the optics and analysis parameters in SMLM, a larger cell diameter results in images with less DNA signal being projected into the central region of the 2D cross-section. In RID plots, DNA signal thus appears closer to the cell boundaries, while cells with regular rod-shape provide RID plots with a broader intensity distribution (see **Figures 4** and **5** as examples). Despite its sectioning capabilities, confocal imaging is also highly affected by 2D projection. Assuming an emission wavelength of 600 nm, the use of a 1.4 NA oil objective and a pinhole size of 1 Airy unit, the axial resolution of a confocal image would amount ~ 600 nm. As this projection volume is larger than the respective volume in SMLM, nucleoids appear less membrane-associated in confocal images and the resulting population averages (see **Figure 3**).

##### Supplementary Note 2: Radial intensity distribution (RID)

The radial intensity distribution (RID) describes how a signal is spatially distributed from an objects' boundary (here the membrane) towards its centroid. The RID is determined by successively eroding the object boundaries and measuring the intensity in the resulting area. The intensity in the eroded layer can be easily calculated by subtracting signal intensities measured before and after the erosion step. Normalization to the total signal intensity provides the relative intensity within the eroded layer (see **equation 3** in the main text). As individual cells have different cellular dimensions, RID analysis requires a variable number of erosion steps. The number of erosion steps or absolute cell width can thus not be used as x-coordinate in RID plots. To circumvent this limitation, we measured the object area before each erosion step and normalized it to the original object area. As objects are eroded radially, the relative cell area correlates with cell width and can be used as approximation for the distance to the cell centre.

#### Supplementary Figures

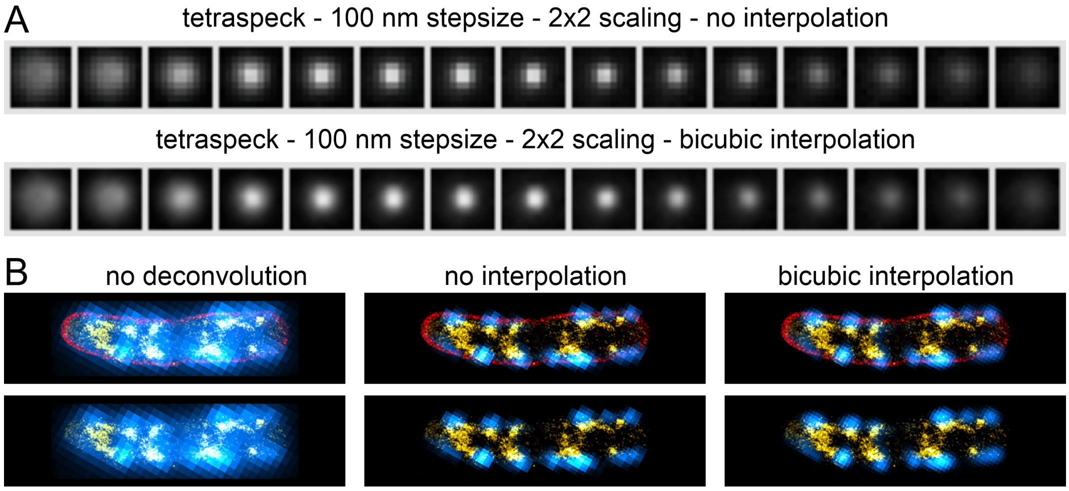


*Figure S1: Deconvolution of MreB-sfGFP$^{sw}$ signal. [A] Representative stack of an individual bead (100 nm tetraspeck), 2x scaled without and with interpolation (bicubic). [B] Exemplary E. coli NO34 cell showing the effect of deconvolution using the Fiji plugin "3D iterative deconvolution" (see methods). 3-4 center planes of the MreB-sfGFP$^{sw}$ stack (cyan hot) were averaged and overlaid with the DNA (yellow hot) and membrane (red) PAINT signal. Deconvolution clearly enhances resolution and contrast of the MreB signal.*

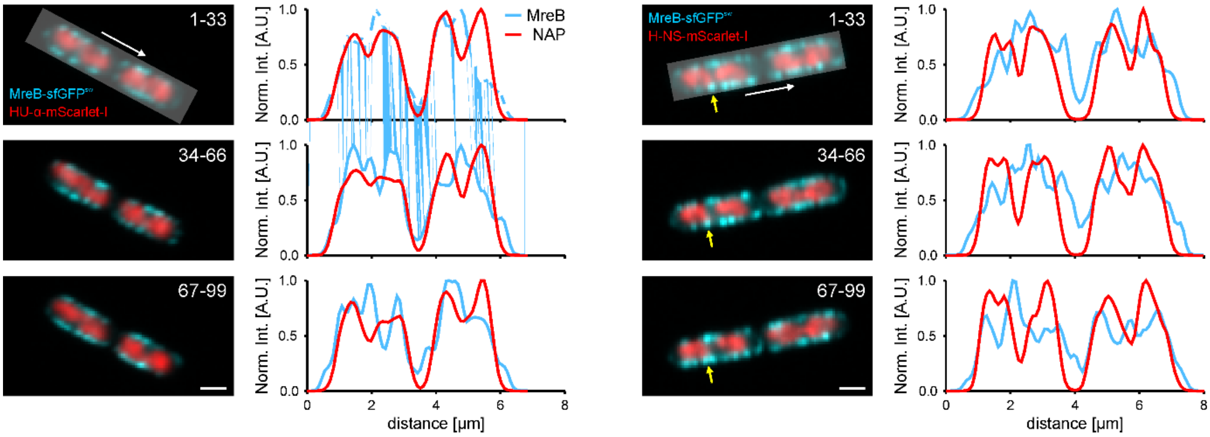


*Figure S2: Representative confocal images of live E. coli cells expressing MreB-sfGFP (cyan) and HU-α-mScarlet-I (left panel, red) or H-NS-mScarlet-I (right panel, red). A subset of 33 frames were averaged and intensities were plotted along the bacterial long axis (white arrow and shaded area). Yellow arrow highlights an elongasome that stays close to a nucleoid filament over the entire time course of imaging. Scale bar is 1 µm.*

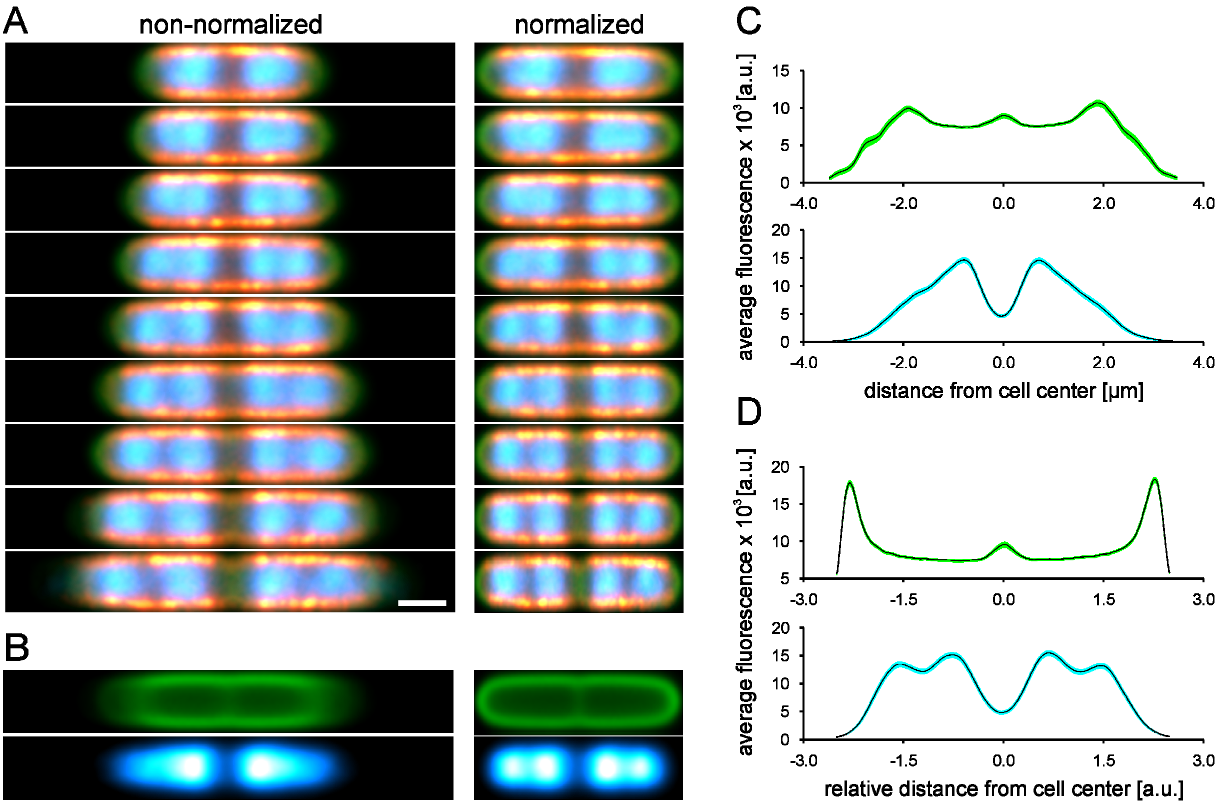


*Figure S3: Averaging of non-normalized and normalized confocal images. [A] Average images of individual E. coli NO34 cells of varying length. 40 cells were averaged for each image with 20 cells overlap. MreB-sfGFP$^{sw}$ is shown in red hot, membrane in green and the nucleoid in cyan hot. Color-code was changed for visualization purpose. [B] Average images of all cells (membrane and nucleoid), either non-normalized (left) or normalized (right). [C] Intensity plot along the length axis of non-normalized average image shown in [B]. [D] Intensity plot along the length axis of normalized average image shown in [B]. The bilobed structure of the sister chromosomes is visible in contrast to the plot in [C]. Scale bar in [A] is 1 µm.*

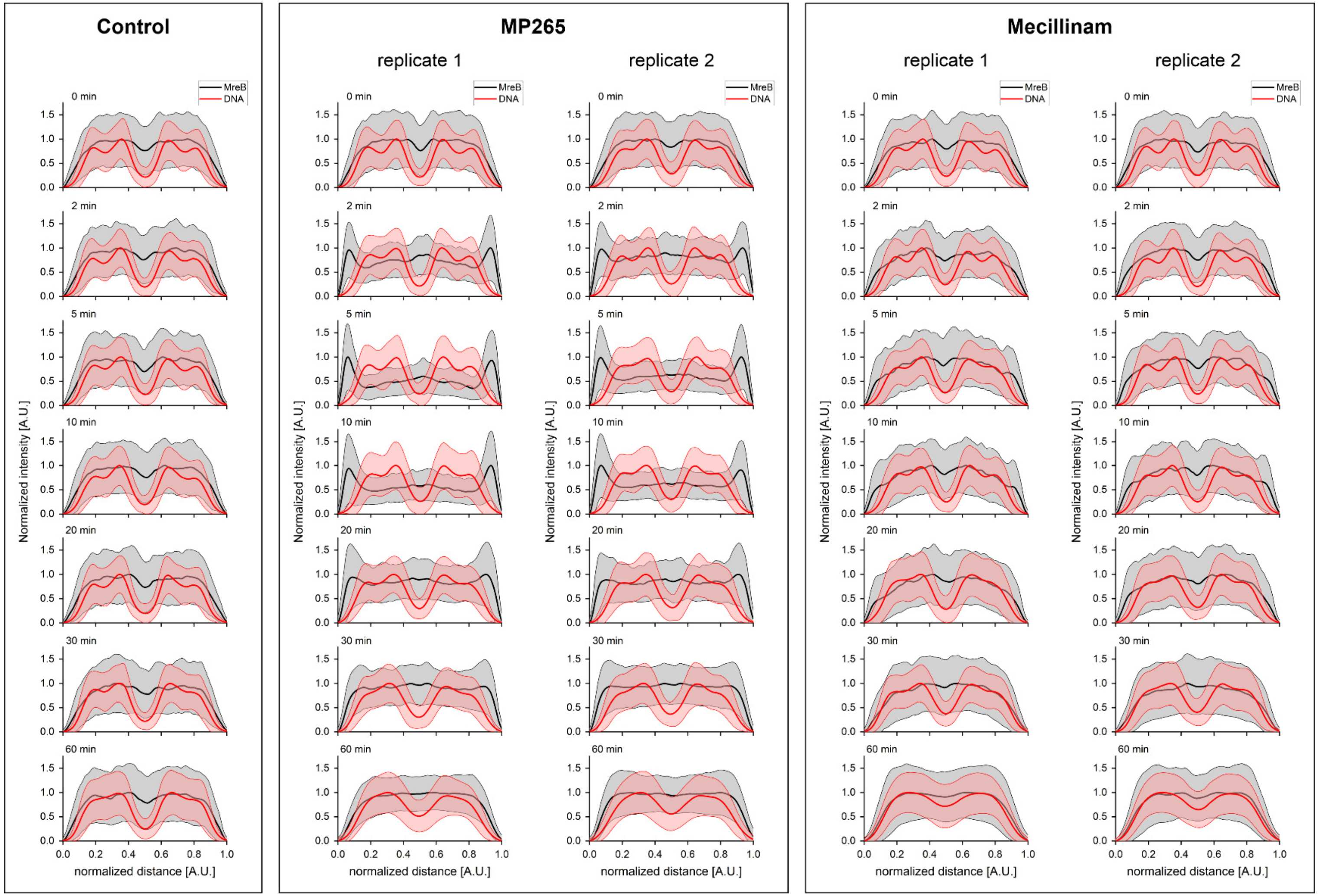


*Figure S4: Length axis plots of MreB and DNA signal during perturbation of cell wall synthesis. Intensities were measured in confocal average and standard deviation images. Lines indicate the mean intensities and shaded areas the standard deviation.*

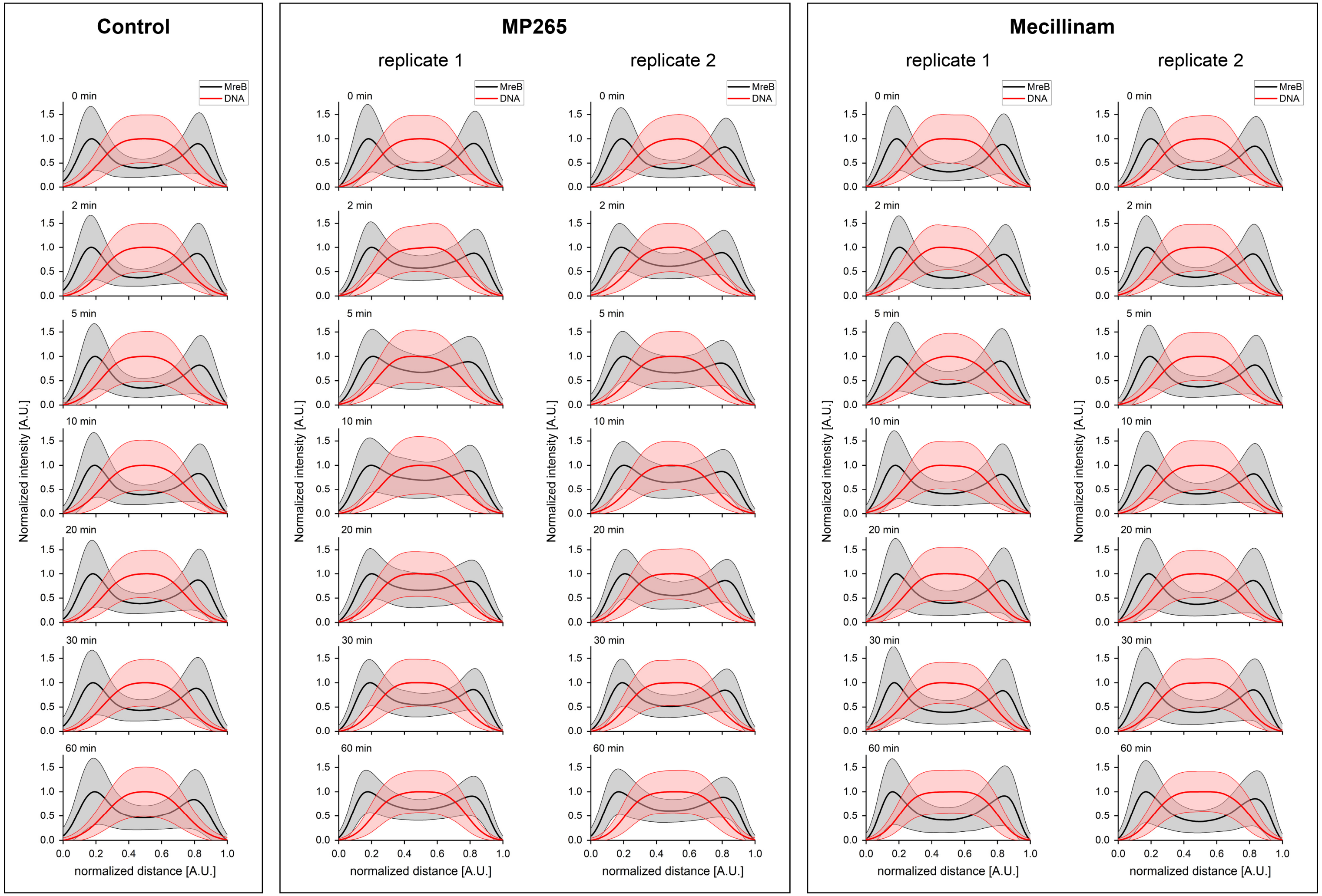


*Figure S5: Cross axis plots of MreB and DNA signal during perturbation of cell wall synthesis. Intensities were measured in confocal average and standard deviation images. Lines indicate the mean intensities and shaded areas the standard deviation.*

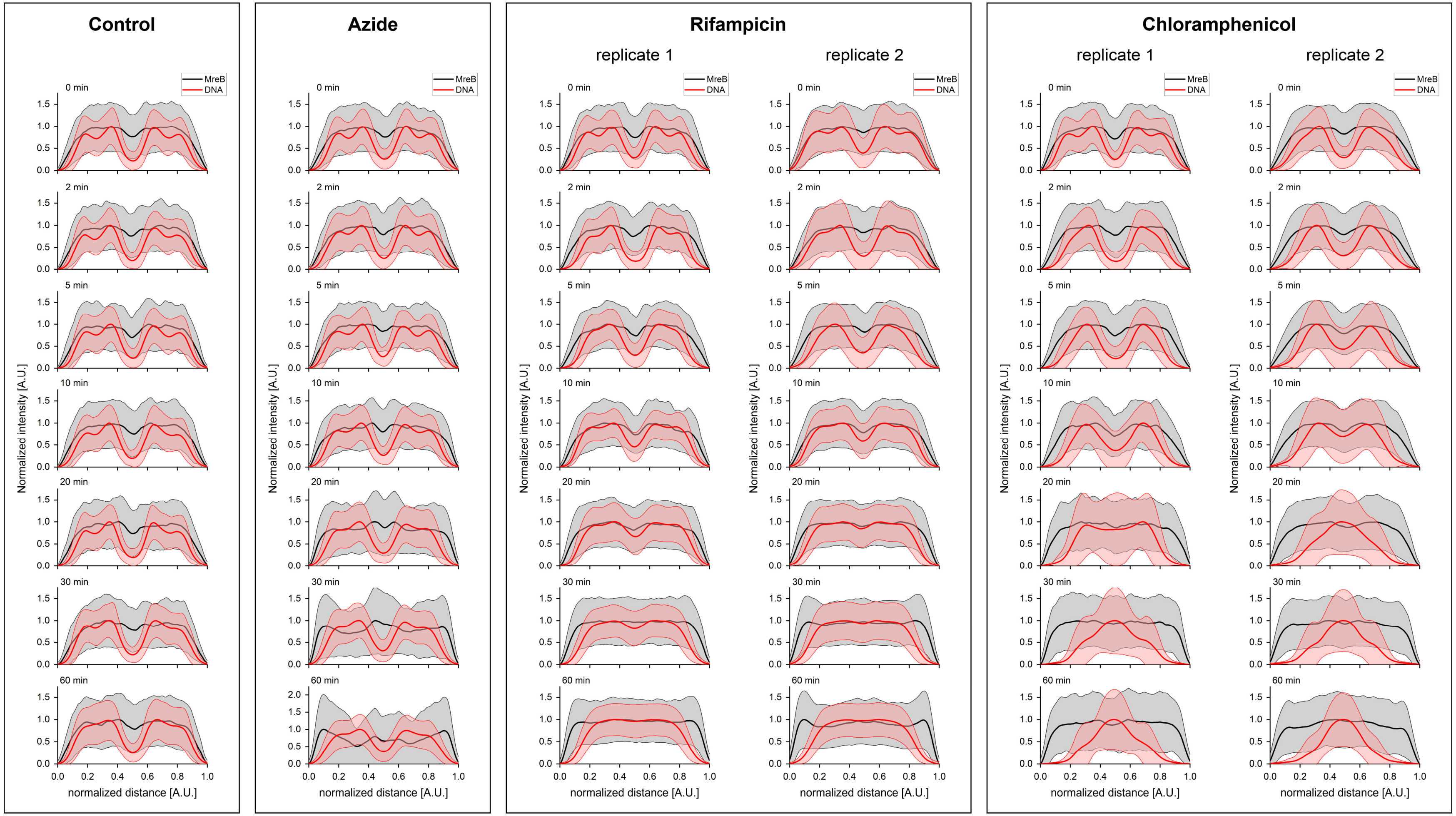


*Figure S6: Length axis plots of MreB and DNA signal during perturbation of protein biosynthesis and transport. Intensities were measured in confocal average and standard deviation images. Lines indicate the mean intensities and shaded areas the standard deviation.*

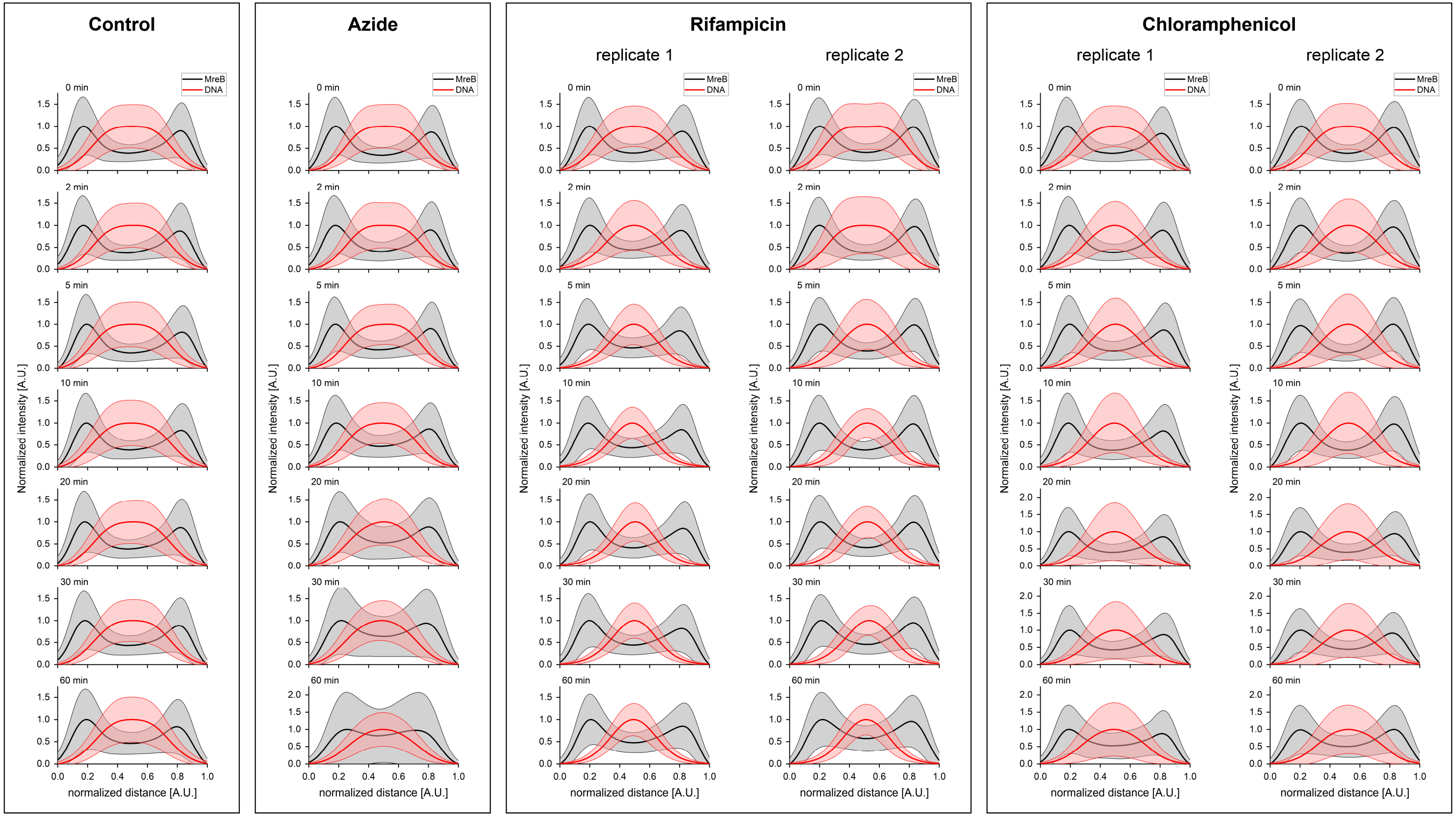


*Figure S7: Cross axis plots of MreB and DNA signal during perturbation of protein biosynthesis and transport. Intensities were measured in confocal average and standard deviation images. Lines indicate the mean intensities and shaded areas the standard deviation.*

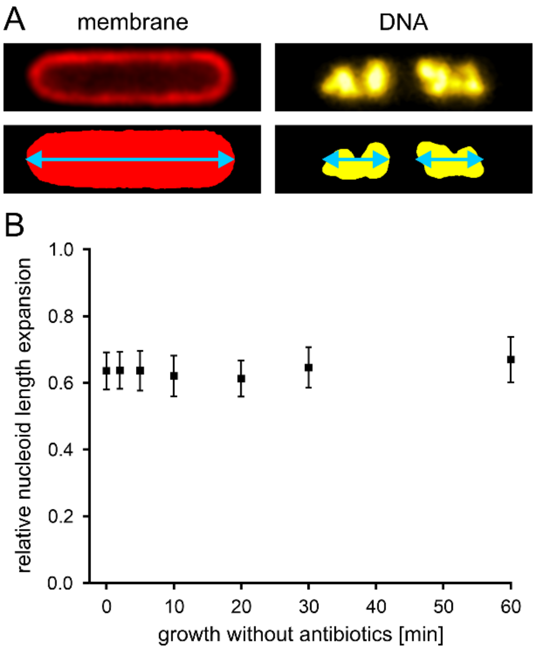


*Figure S8: Measurement of relative nucleoid length of untreated cells over time. [A] Representative NO34 cell labeled for membrane (Nile red, red) and DNA (DAPI, yellow hot). Signal is binarized and cell (left) and nucleoid (right) lengths are measured using a custom-written Fiji macro. [B] Relative nucleoid length over time. Aliquots of a control culture were fixed at the indicated time points and imaged using confocal microscopy. The average relative nucleoid length does not change with increasing culture density during the observation period. Data points represent mean values and error bars the standard deviation.*

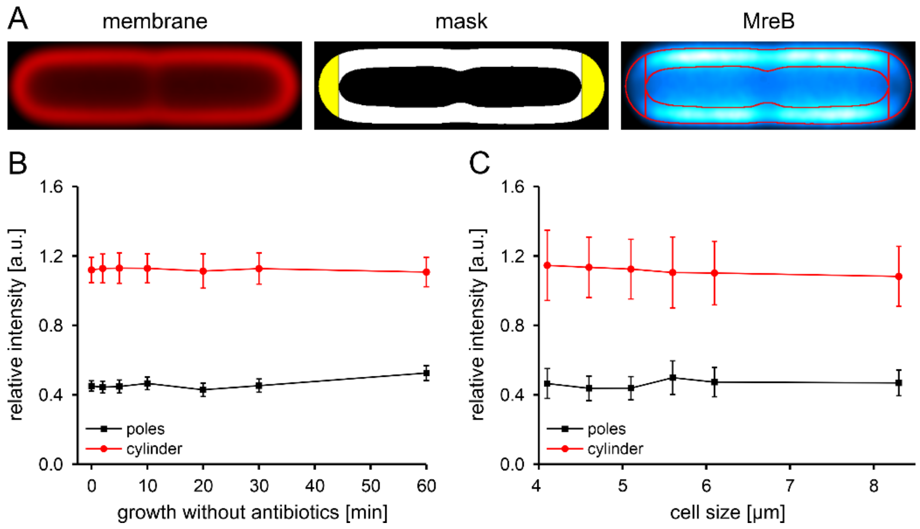


*Figure S9: Analysis of the MreB intensity distribution along the bacterial membrane in confocal average images. [A] Schematic of the analysis. The average membrane image is thresholded using Otsu's method. Poles (mid panel, yellow areas) are segmented according to the cytosol (black area). Pole (yellow areas) and cylindrical membrane sections (white areas) are used to measure the MreB-sfGFP$^{sw}$ intensity in the MreB average image (right panel). [B] Relative MreB intensity over time in an untreated culture. MreB distribution remains constant over the entire time course. [B] Length-dependence of the MreB distribution. A dataset (untreated culture, t = 0 min) was split into subsets according to cell length (0.5 µm interval except > 6 µm) and average images were calculated. MreB intensity distribution is insensitive to cell length. Higher errors are a result of the lower cell count used for generation of the average images. Data points represent individual values and error bars the standard error of the mean calculated from standard deviation images of the averaging process.*

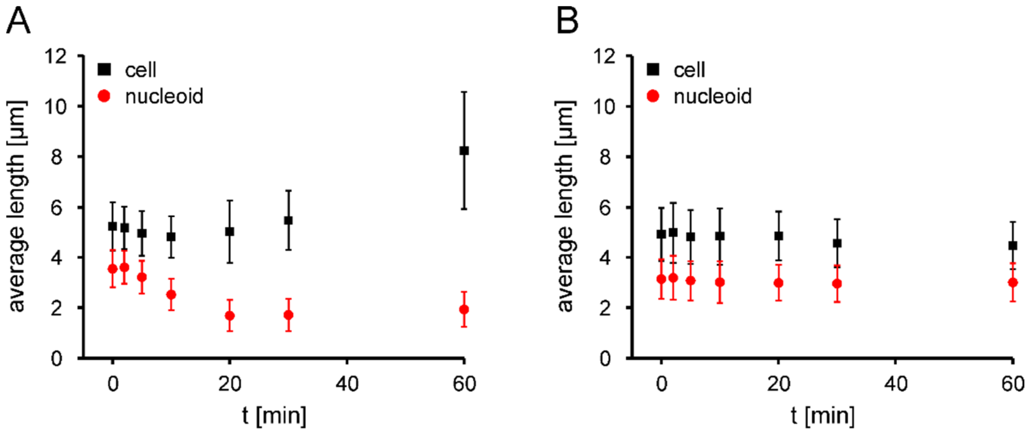


*Figure S10: Measurement of cell and nucleoid lengths during nalidixate treatment. [A] Average length over time during exposure to nalidixate. Cell length (black) increases with nalidixate exposure, while nucleoid length (red) shrinks until it reaches a plateau after 20 min. [B] Average length over time in an untreated control culture. Average cell and nucleoid lengths remain constant during the observation period. Data points represent mean values and error bars the respective standard deviation.*

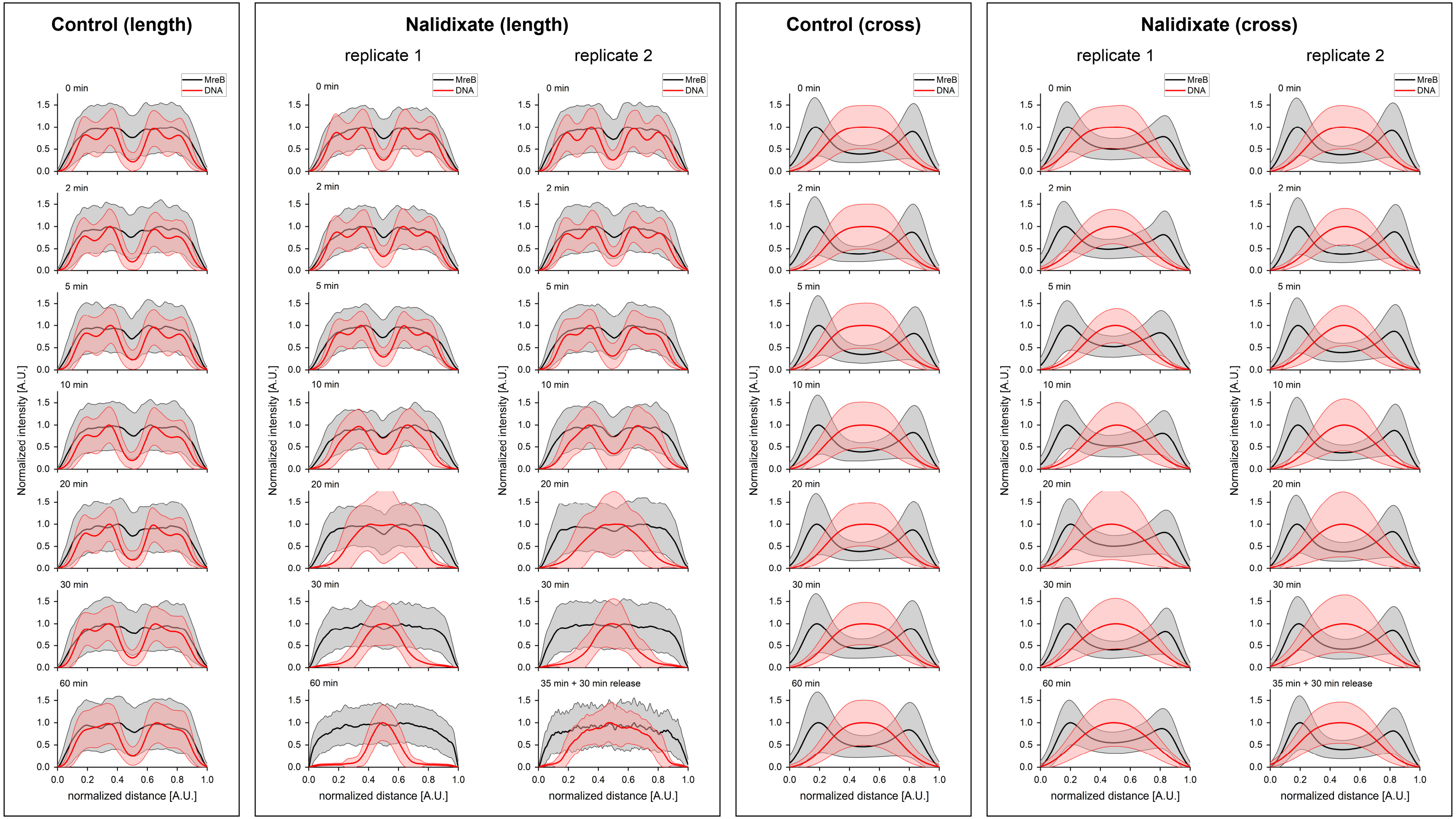


*Figure S11: Length and cross axis plots of MreB and DNA signal inhibition of DNA replication. Intensities were measured in confocal average and standard deviation images. Lines indicate the mean intensities and shaded areas the standard deviation.*

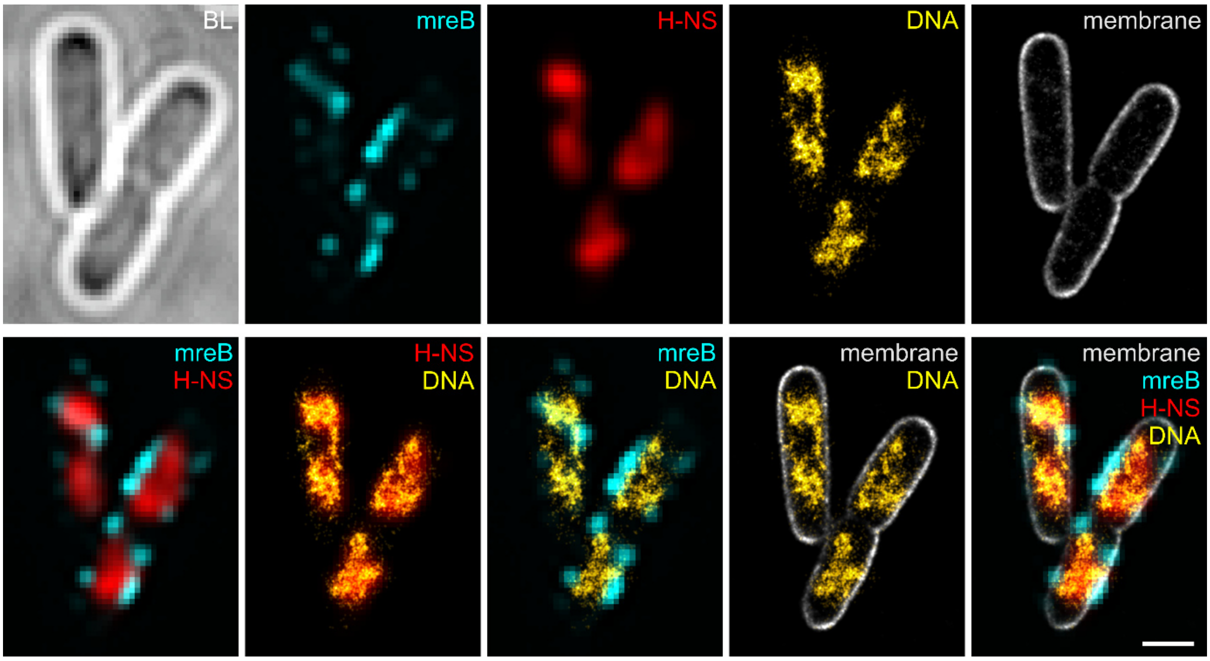


*Figure S12: Dual-color PAINT imaging of fixed E. coli expressing MreB-sfGFP$^{sw}$ and H-NS-mScarlet-I from the native locus. The images were acquired on a commercial N-STORM setup. Upper row shows the individual channels, while the lower row shows selected overlays. The super-resolution PAINT image matches the signal obtained from H-NS-mScarlet-I. Scale bar is 1 µm.*

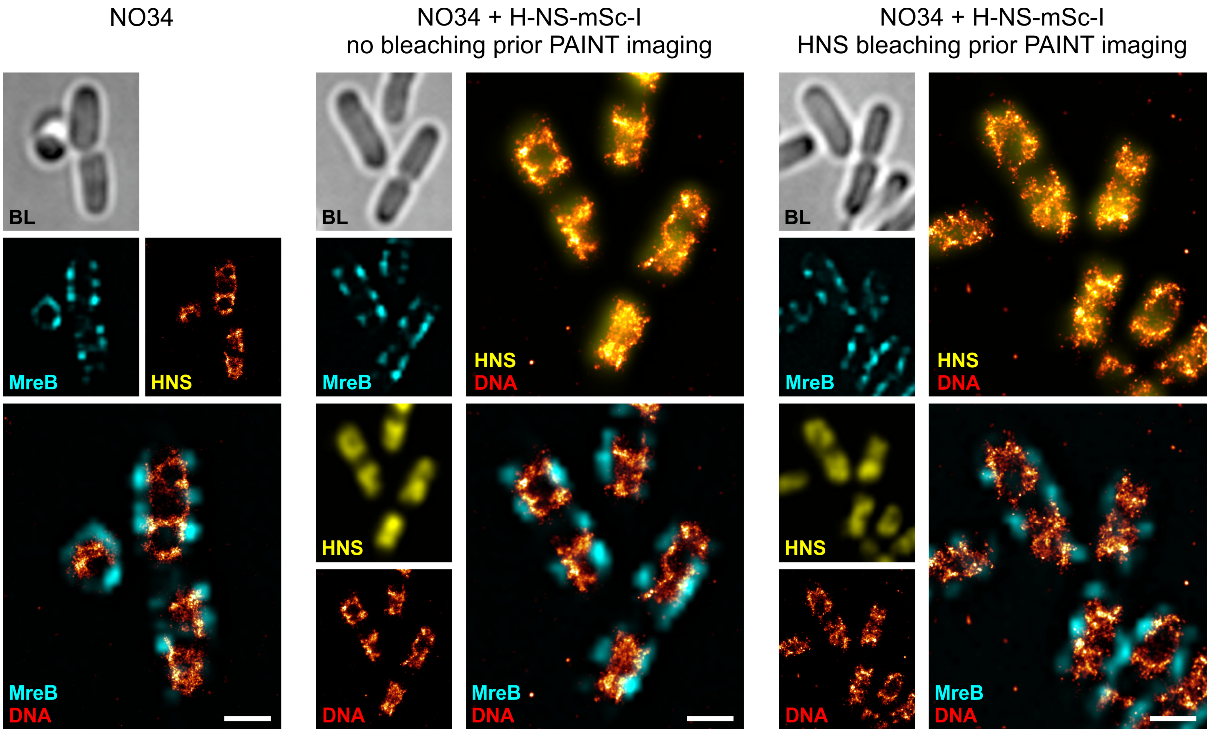


*Figure S13: PAINT imaging of the nucleoid in fixed E. coli cells expressing MreB-sfGFP$^{sw}$ and H-NS-mScarlet-I from the native locus. Images were acquired on a commercial Elyra PS1 setup. The super-resolution PAINT image matches the signal obtained from H-NS-mScarlet-I. Prebleaching H-NS-mScarlet-I was tested to reduce the background signal in the red channel (JF$_{646}$-Hoechst, DNA), which had no significant effect on the PAINT image quality. Scale bar is 1 µm.*

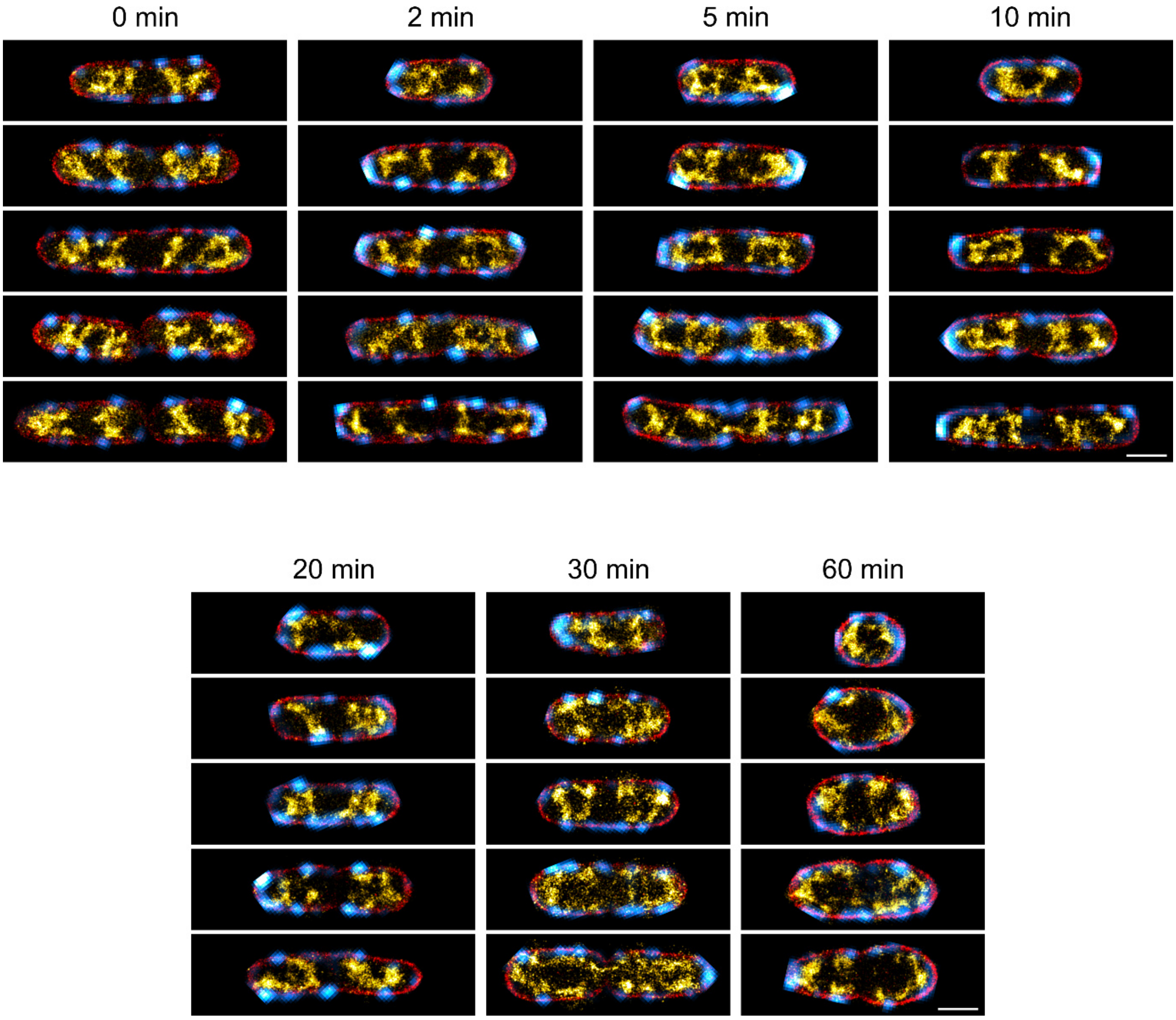


*Figure S14: Exemplary dual-color PAINT images of fixed E. coli NO34 cells treated with 25 µM MP265. MreB-sfGFP$^{sw}$ is shown in cyan hot, membrane in red and DNA in yellow hot. Scale bars are 1 µm.*

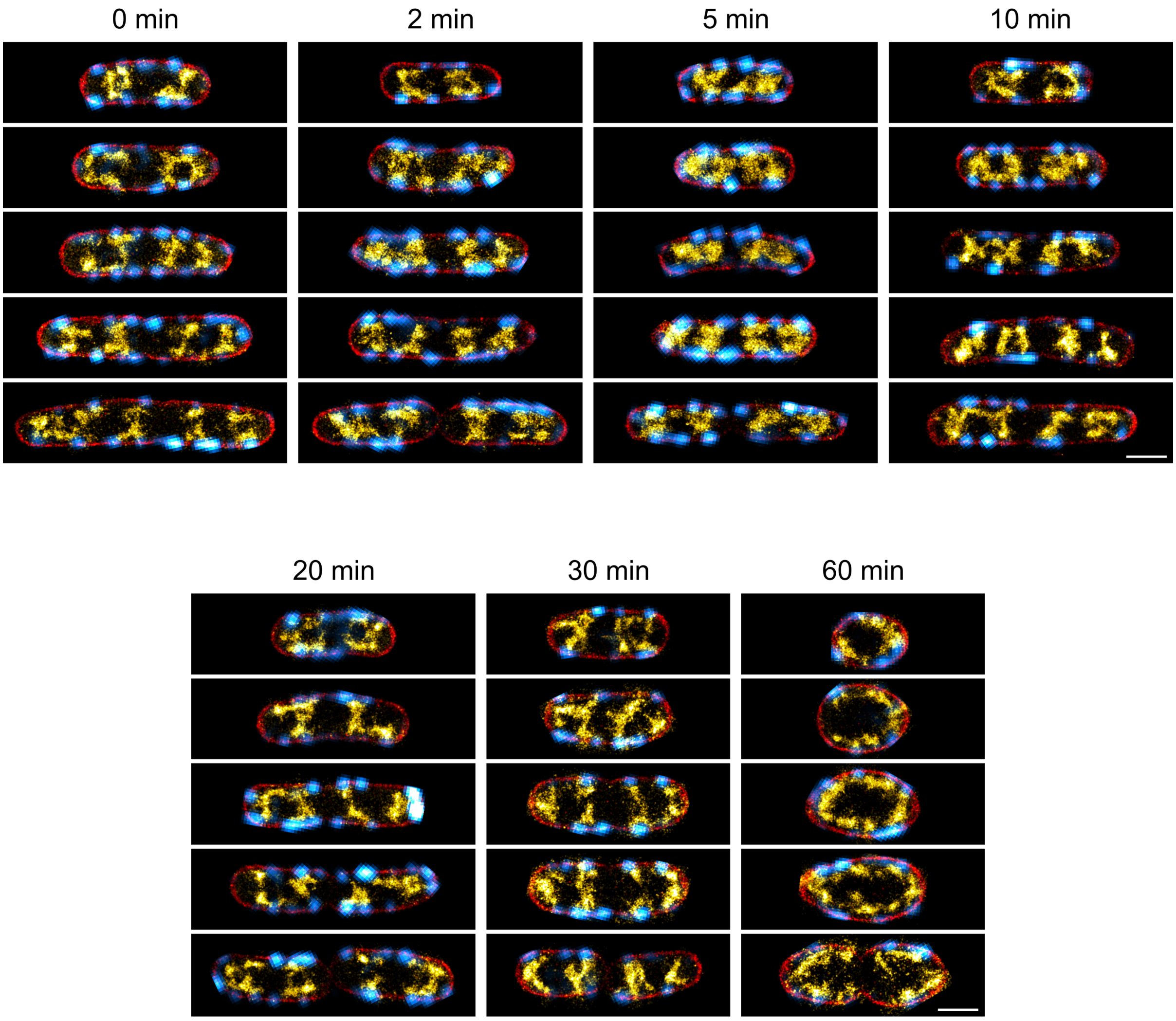


*Figure S15: Exemplary dual-color PAINT images of fixed E. coli NO34 cells treated with 2 µg/ml Mecillinam. MreB-sfGFP$^{sw}$ is shown in cyan hot, membrane in red and DNA in yellow hot. Scale bars are 1 µm.*

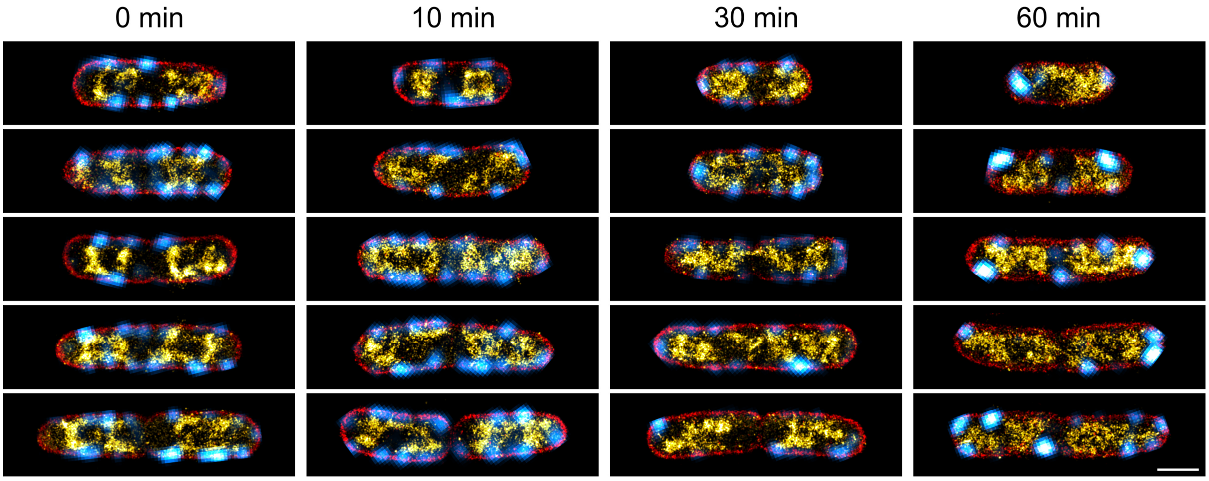


*Figure S16: Exemplary dual-color PAINT images of fixed E. coli NO34 cells treated with 1 µM sodium azide ($NaN_3$). MreB-sfGFP$^{sw}$ is shown in cyan hot, membrane in red and DNA in yellow hot. Scale bars are 1 µm.*

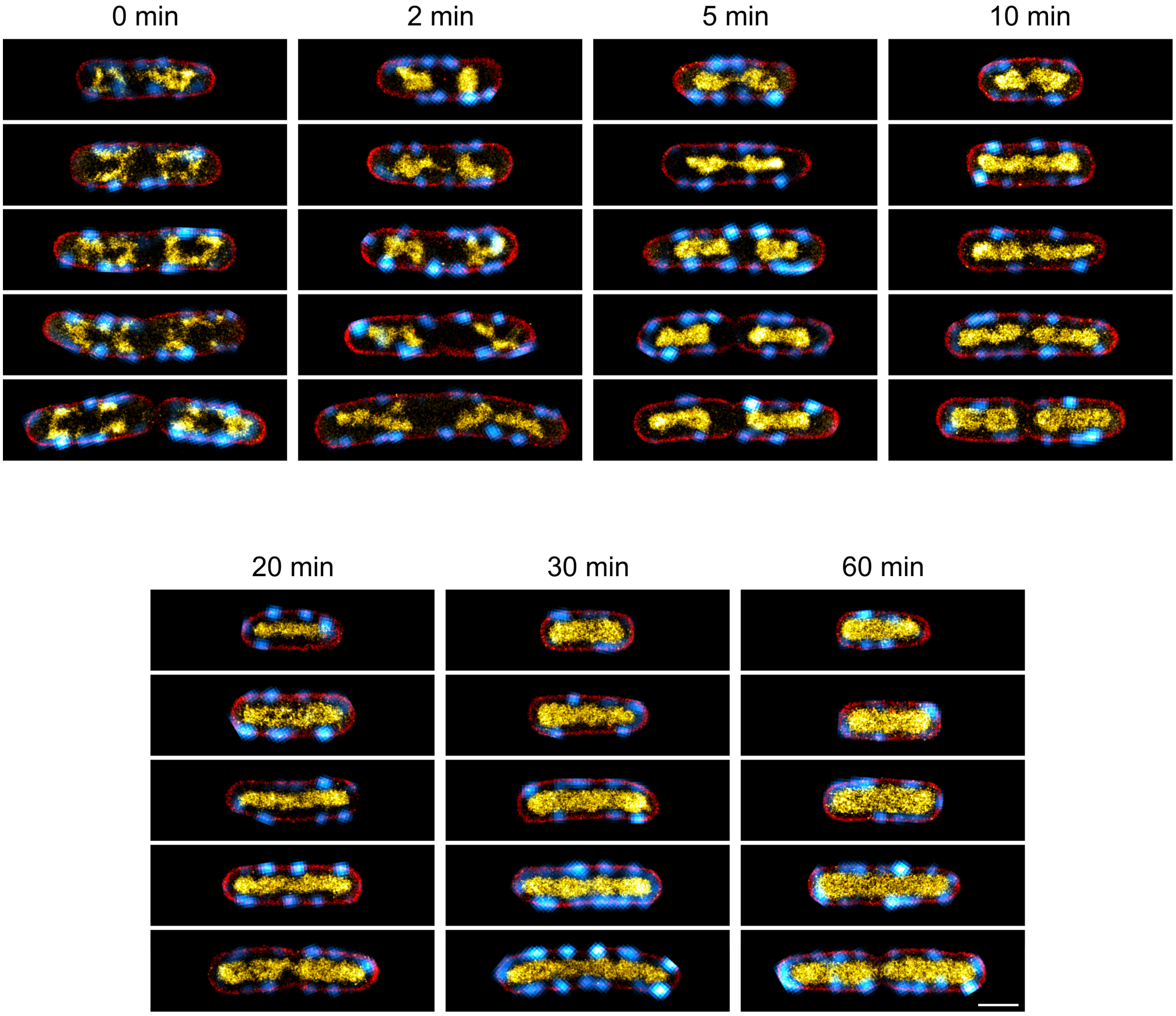


*Figure S17: Exemplary dual-color PAINT images of fixed E. coli NO34 cells treated with 100 µg/ml rifampicin. MreB-sfGFP$^{sw}$ is shown in cyan hot, membrane in red and DNA in yellow hot. Scale bars are 1 µm.*

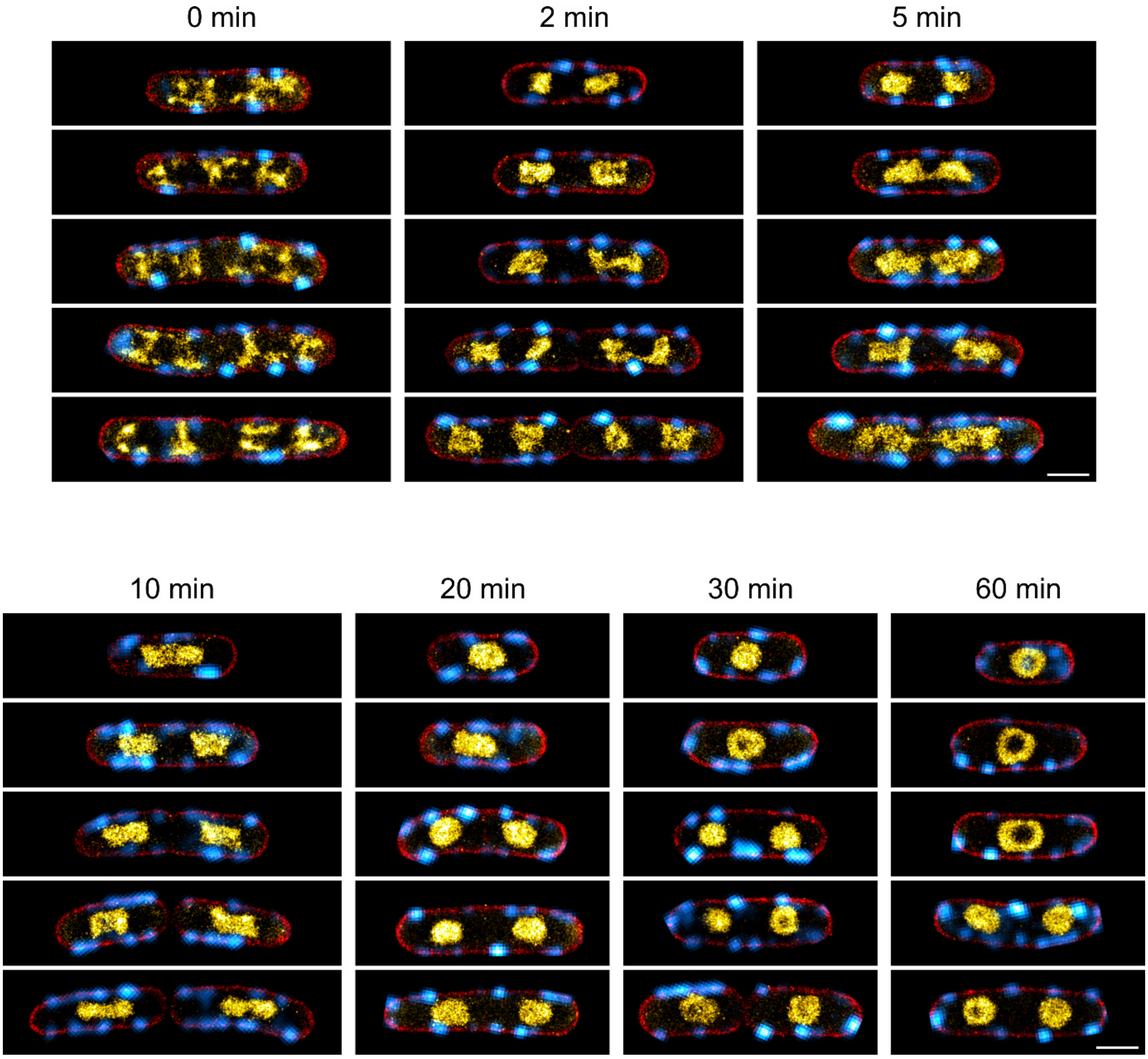


*Figure S18: Exemplary dual-color PAINT images of fixed E. coli NO34 cells treated with 50 µg/ml chloramphenicol. MreB-sfGFP$^{sw}$ is shown in cyan hot, membrane in red and DNA in yellow hot. Scale bars are 1 µm.*

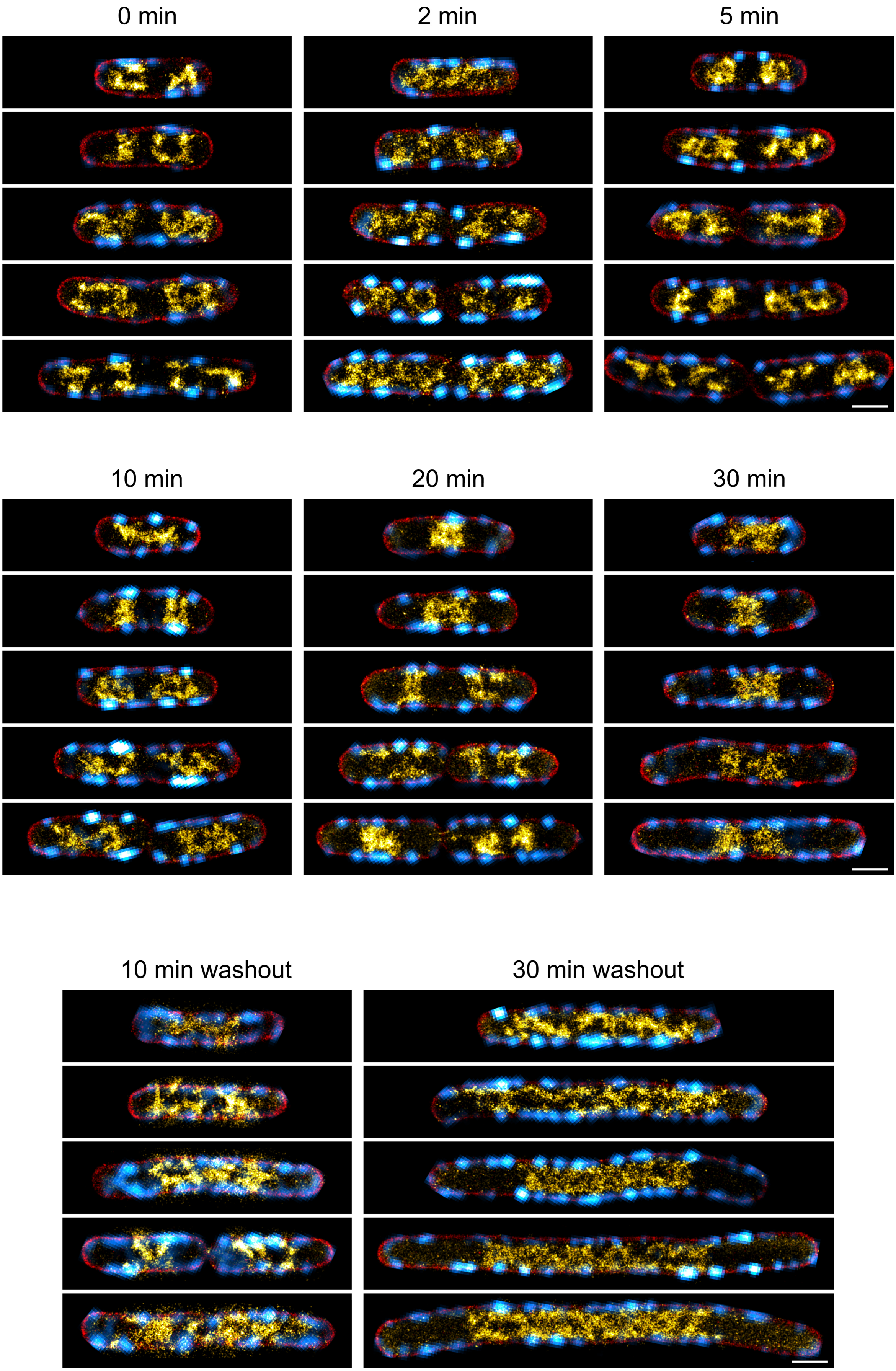


*Figure S19: Exemplary dual-color PAINT images of fixed E. coli NO34 cells treated with 100 µg/ml nalidixate. MreB-sfGFP$^{sw}$ is shown in cyan hot, membrane in red and DNA in yellow hot. Scale bars are 1 µm.*

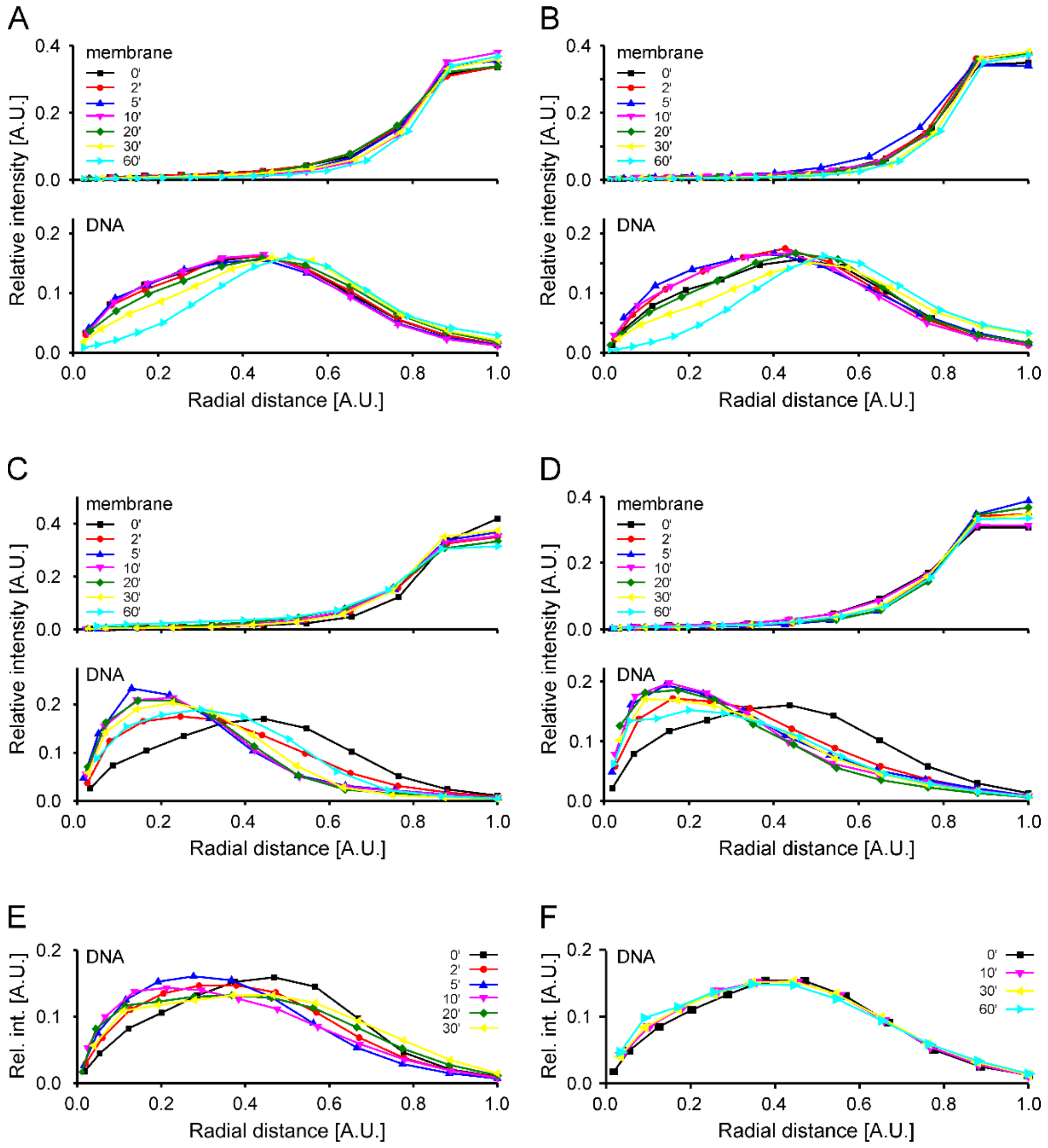


*Figure S20: Erosion analysis plots of membrane and DNA signals for different drug treatments and time points. [A] MP265 treatment. [B] Mecillinam treatment. [C] Rifampicin treatment. [D] Chloramphenicol treatment. [E] DNA signal of cells treated with nalidixate. [F] DNA signal of cells treated with sodium azide. Values represent mean values. Error bars are omitted for visualization.*

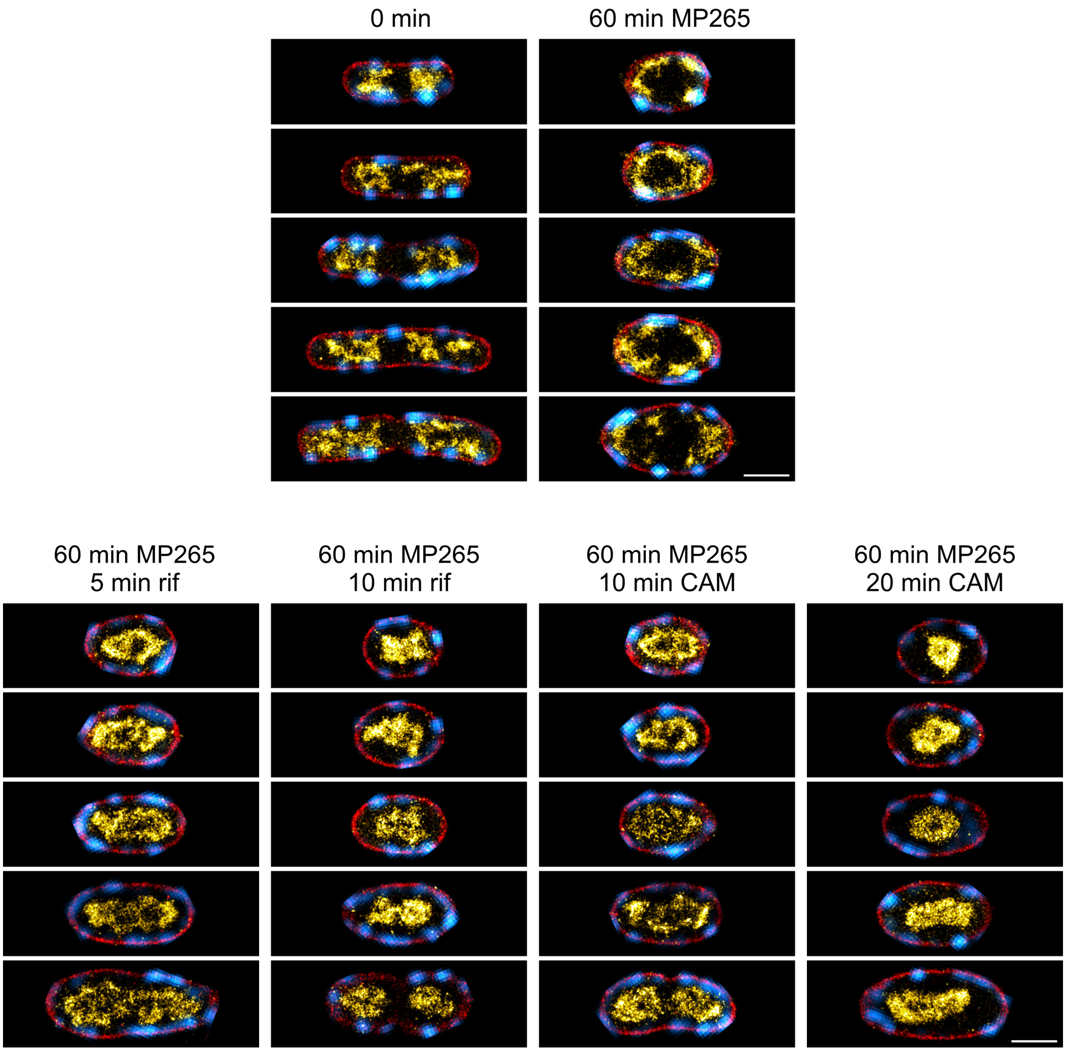


*Figure S 21: Exemplary dual-color PAINT images of fixed E. coli NO34 cells treated with 25 µM MP265 and 100 µg/ml rifampicin or 50 µg/ml chloramphenicol. Scale bar is 1 µm.*

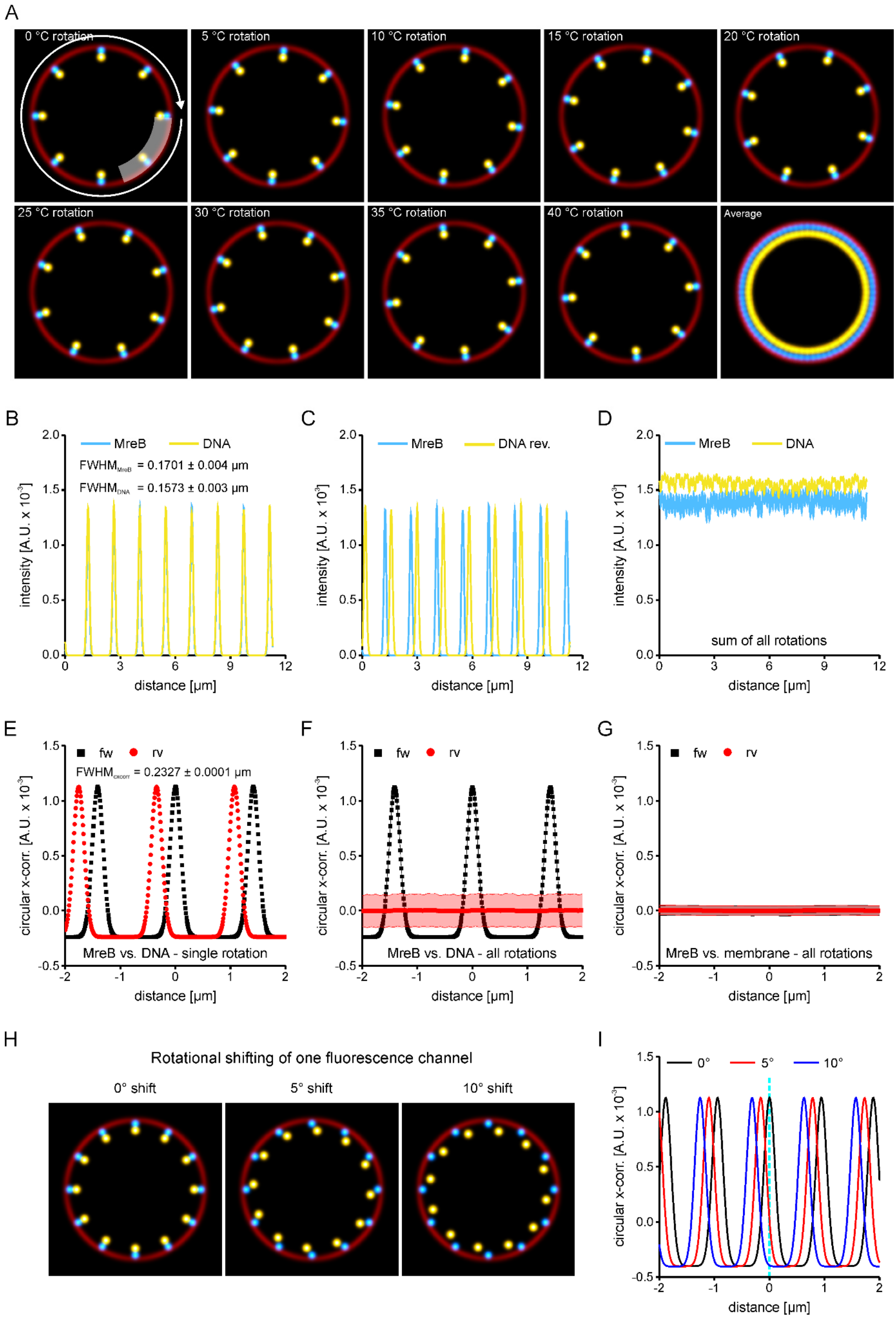


*Figure S22: Circular cross correlation of simulated data. [A] 8 equidistantly spaced spots were positioned on a circle (mimicking the bacterial membranes) with displacement towards the center to mimic MreB and DNA. Heterogeneity is introduced by*

*rotating the spots with an increment of 5°. The average image shows the complete coverage of the circle area. The intensity trace is measured by a line plot of specific thickness along the circle perimeter (white arrow and shaded area). [B] Intensity trace for a single rotation in forward direction (same direction of MreB and DNA traces). [C] Intensity trace for a single rotation in reverse direction (DNA trace was inverted). The signal is shifted as the circular selection is opened at an arbitrary position. Thus, the inverted trace is shifted based on the starting point of the line plot. [D] Intensity traces of the average image shown in [A]. [E] Circular cross-correlation (x-corr) of a single rotation. For intensity traces aligned in the same direction, a peak is observed at the zero position. Neighboring peaks represent correlations to neighboring spots. The FWHM of 0.2327 hereby matches the convolution of the FWHM of both signals as shown in [B]. [F] Average circular x-corr of all rotations shown in [A]. The cross-correlations of the original intensity traces do not change, while the they average out for MreB signals and the inverted DNA intensity traces. [G] Control analysis of the MreB signal vs the membrane signal. In both original and reverse mode, the analysis expectedly shows no correlation. [H] Simulation of images with shifted channels. DNA signal was shifted by 5° or 10°. [I] Circular cross-correlations of the images shown in H. The shift of the DNA channel leads to a shifted peak in the cross-correlation analysis.*

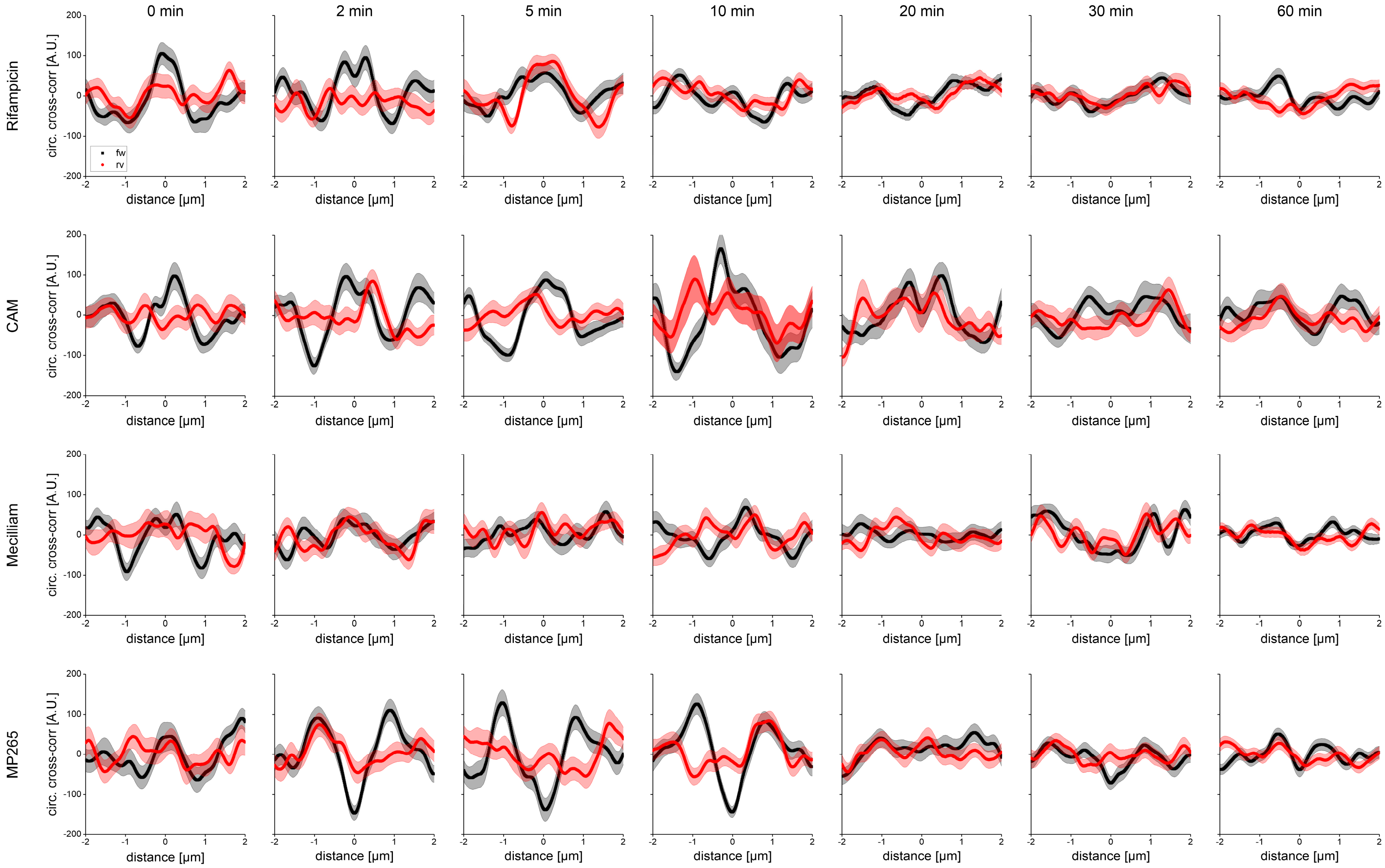


*Figure S23: Circular x-corr analysis of drug treatments. Correlations of original traces are shown in black, while correlation of MreB with inverted DNA signals is shown in red. Lines represent mean values und shaded areas the standard error of the mean.*

#### Supplementary Tables

*Table S1: Bacterial strains used in this study*

| Bacterial strain | Genotype | Reference/source |
|---|---|---|
| MG1655 | *Escherichia coli* K-12 wild type | CGSG #6300 |
| NO34 | MG1655 *mreB$^{sw}$-sfGFP csrD::*kan | Gitai laboratory [1] |
| CS1 | MG1655 *mreB$^{sw}$-sfGFP csrD::*kan *hupA::mScarlet-I-frt-cam-frt* | this work |
| CS2 | MG1655 *mreB$^{sw}$-sfGFP csrD::*kan *hns::mScarlet-I-frt-cam-frt* | this work |
| CS_Xd1 | *Xenorhabdus doucetiae* DSM17909 *mreB$^{sw}$-sfGFP* | this work |

*Table S2: Primers used in this study*

| Primer name | Sequence (5' – 3') |
|---|---|
| CS_FFM_002 | CAACTGGTTGGTTTCGGTACC |
| CS_FFM_003 | GAATTCACCAGAACCAGCAGCAGAACCAGCAGAACCCTTAACTGCGTCTTTCAGTGCC |
| CS_FFM_005 | TGCTGCTGGTTCTGGTGAATTCGTGAGCAAGGGCGAGGC |
| CS_FFM_006 | TTACTTGTACAGCTCGTCCATGCC |
| CS_FFM_012 | AAGCTAAACGTGCTCAGCGTC |
| CS_FFM_013 | GAATTCACCAGAACCAGCAGCAGAACCAGCAGAACCTTGCTTGATCAGGAAATCGTCG |
| CS_FFM_017 | GGCATGGACGAGCTGTACAAGTAAGCATGGATGAGCTGTACAAATAAG |
| CS_FFM_018 | CAGAAAGACAAAAGGGGTGAAACCACCCCTTCGTTAAAACTGTTCACTGCCACGCAAT CATTGTAGGCTGGAGCTGCTTC |
| CS_FFM_019 | CAGAAAGACAAAAGGGGTGAAAC |
| CS_FFM_020 | CAATAAAAAATCCCGCCGCTGGCGGGATTTTAAGCAAGTGCAATCTACAAAAGAATTG TAGGCTGGAGCTGCTTC |
| CS_FFM_021 | CAATAAAAAATCCCGCCGCTGG |
| CS_MPI_003 | ATCGATCCTCTAGAGTCGACCTGCACATCACCACAATTTTTCATCAC |
| CS_MPI_004 | CGTGGGATAGGCAGAAC |
| CS_MPI_005 | GATGAAGTACGGGAAATTGAAG |
| CS_MPI_006 | TGGAATTCCCGGGAGAGCTCAGATCTTAACGAATTGAAAATTGACGACG |
| CS_MPI_018 | TGAAATTGGTTCTGCCTATCCCACGTCTGGCTCGAGCAGTAAAGG |
| CS_MPI_019 | GCACTTCAATTTCCCGTACTTCATC |
| CS_MPI_020 | GAAGCTATCATCAATTATGTTCGC |
| CS_MPI_021 | TGCTTCAAGAATTTCATTGGAATTCAG |
| VpDS132 fw | GATCGATCCTCTAGAGTCGACC |
| VpDS132 rv | CACATGTGGAATTGTGAGCGG |

*Table S3: Macros and notebooks used in this study*

| Macro | Purpose |
|---|---|
| M1 | Average PSF determination from bead stacks |
| M2 | Smoothing of cell outlines in NR PAINT images |
| M3 | Rotation, alignment and cell straightening – CLSM images |
| M4 | Cell normalization – CLSM images |
| M5 | Determination of the relative nucleoid length expansion – CLSM images |
| M6 | Determination of the relative MreB distribution – CLSM images |
| M7 | Determination of radial intensity distributions using erosion analysis |
| M8 | Plot intensity profiles along perimeter |
| M9 | Simulate images for circular cross-correlation |
| Notebook_1 | Google Colab notebook for the calculation of circular cross-correlation |

*Table S4: Identifier of datasets made publicly available via Zenodo*

| Dataset | DOI |
|---|---|
| CLSM images of all treatments - *Escherichia coli* | 10.5281/zenodo.8430052 |
| Nucleoid segmentation SMLM data | 10.5281/zenodo.8429932 |
| SMLM images of all treatments – *Escherichia coli* | 10.5281/zenodo.8430032 |
| SMLM images of *Xenorhabdus doucetiae* | 10.5281/zenodo.10007398 |

#### Supplementary Videos (Stills)

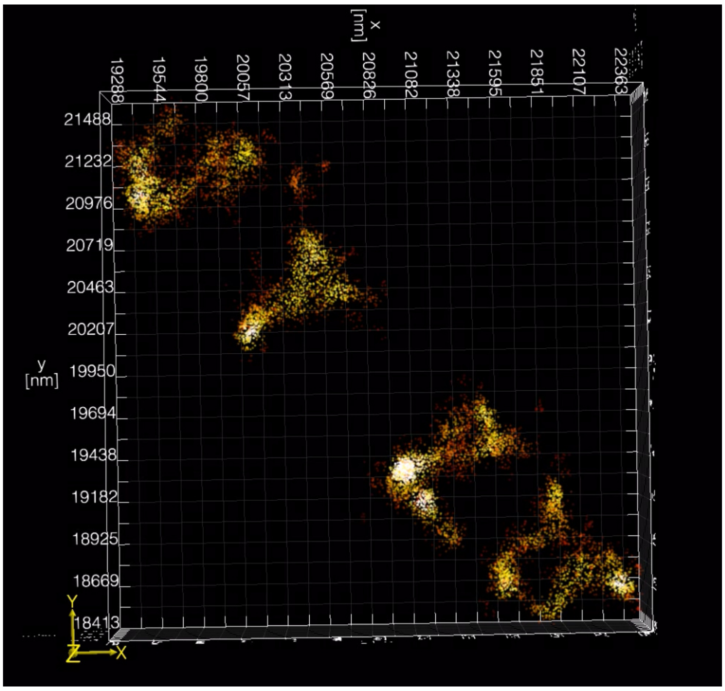


*Supplementary Video 1: Rendering or 3D SMLM imaging of an E. coli cell click-labelled for DNA.*

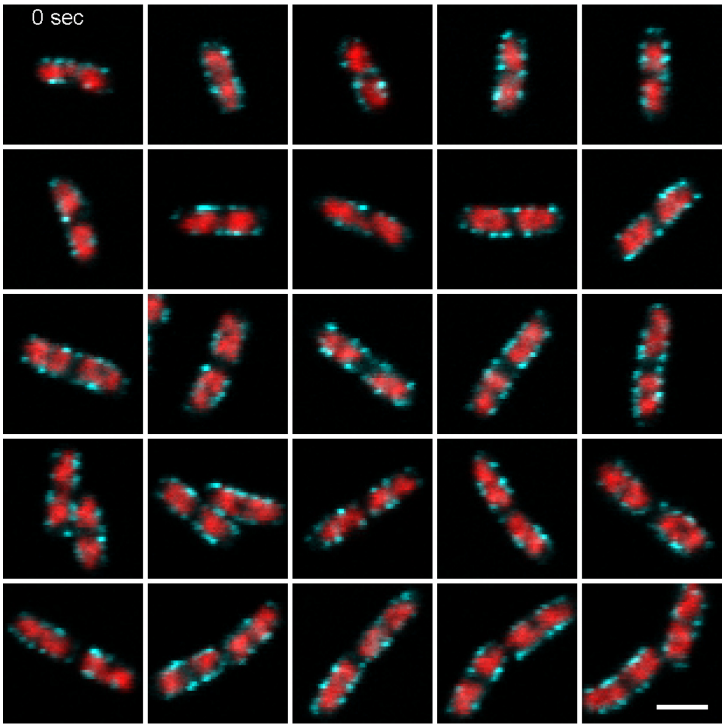


*Supplementary Video 2: Live-cell confocal microscopy time series of E. coli cells labeled for HU-α (HupA-mScarlet-I, red) and MreB (MreB$^{sw}$-sfGFP, cyan). Scale bar is 2 µm.*

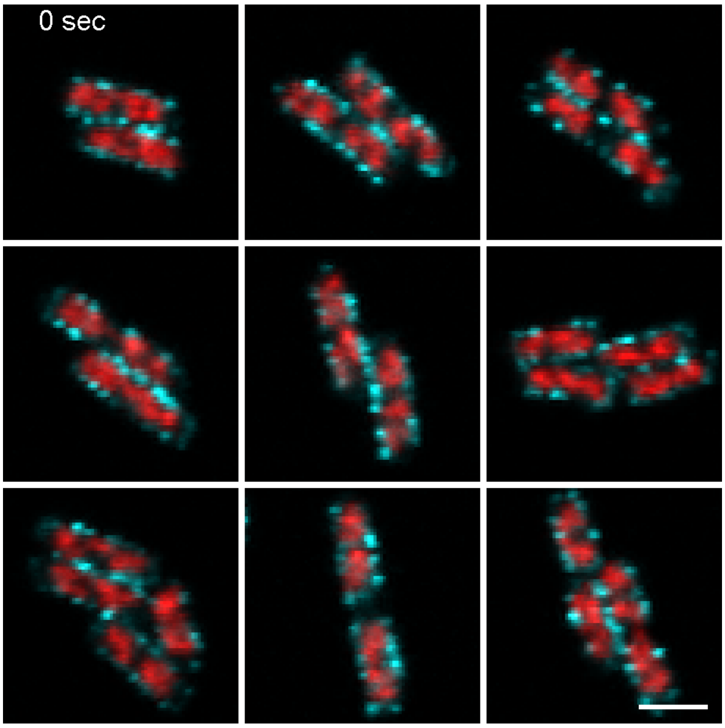


*Supplementary Video 3: Live-cell confocal microscopy time series of E. coli cells labeled for H-NS (H-NS-mScarlet-I, red) and MreB (MreB$^{sw}$-sfGFP, cyan). Scale bar is 2 µm.*

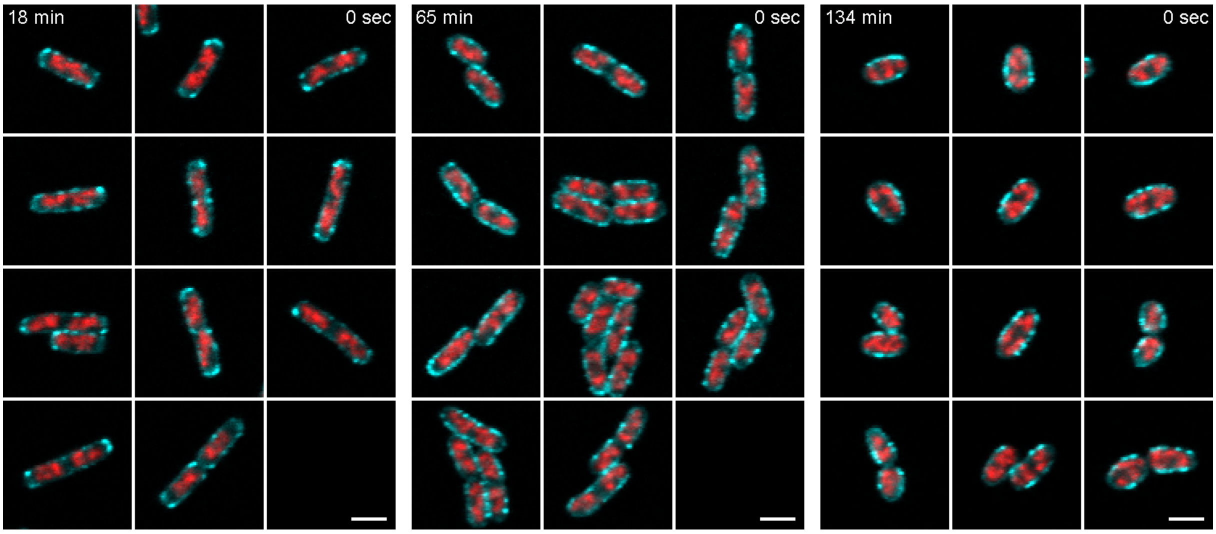


*Supplementary Video 4: Live-cell confocal microscopy time series of E. coli cells labeled for H-NS (H-NS-mScarlet-I, red) and MreB (MreB$^{sw}$-sfGFP, cyan). Cells were imaged under agarose pads containing 25 µM MP265. Time stamps on the left represent the starting point of the measurement after immobilization under the agarose pads, while stamps on the right indicate the time during the time lapse measurement. Scale bar is 2 µm.*
