## Supplementary material for "Transertion and cell geometry organize the *Escherichia coli* nucleoid during rapid growth": Macro_M1 - Average PSF from beadstack.ijm

```
// This macro creates the average PSF from a z-stack of fiducial markers

// This macro is part of the publication: " Transertion and cell geometry organize the Escherichia coli nucleoid during rapid growth "

// Author:	Christoph Spahn
// 		MPI for terrestrial microbiology, Marburg
// 	

// This macro was written in Fiji v1.53t
// It requires the 'GaussFit on Spot' plugin that can be found here:
// https://imagej.nih.gov/ij/plugins/gauss-fit-spot/index.html
// Download the .jar file and place it in the Fiji plugins folder


// ===========================================================
//    This dialog fetches the parameters for PSF extraction

Interpolation_types = newArray("None", "Bilinear", "Bicubic"); // Types of interpolation during image scaling

Dialog.create("Parameters for PSF extraction");
Dialog.addNumber("Scaling factor:", 2); // Oversampling factor to extract the PSF with subpixel accuracy
Dialog.addChoice("Interpolation type:", Interpolation_types, "Bicubic"); // Interpolation type for image scaling
Dialog.addNumber("PSF width:", 13); // Window size for PSF extraction in unscaled pixels (full width)
Dialog.addNumber("Peak prominence:", 500); // Peak prominence for bead detection; determine suitable values using the 'Find Maxima' function
Dialog.addCheckbox("Subtract background", true); // Subtracts the average of slice median values from the entire image stack
Dialog.addCheckbox("Save standard deviation of average PSF", true); // automatically saves the standard deviation image of the averaged PSF; can be useful to check for artifacts
Dialog.show();

scaling_factor = Dialog.getNumber();
interpolation = Dialog.getChoice();
PSF_size = Dialog.getNumber();
peak_prominence = Dialog.getNumber();
subtract_background = Dialog.getCheckbox();
save_PSF_std = Dialog.getCheckbox();
scaled_width = scaling_factor * PSF_size;


// Here, we get the results directory
// A folder for the individual PSFs is generated automatically

run("Clear Results");
roiManager("Reset");

path = getDirectory("Select output directory"); // Results folder

// Here, we create a subfolder in the selected folder to save the individua PSF stacks
path2 = path + "/Single_PSFs/";
File.makeDirectory(path2);

// The image stack should be open and active
stack = getImageID();
run("Scale...", "x=["+scaling_factor+"] y=["+scaling_factor+"] z=1.0 interpolation=["+interpolation+"] average process create");
scaled_stack = getImageID();
getDimensions(width, height, channels, slices, frames); // determines image properties

// In this block, we determine the focal plane of the bead stack
// Typically, the focal plane is the plane with the highest standard deviation as beads have the highest signal

if (frames>slices) {
	slice_count = frames;
}
	else slice_count = slices;

std_statistics = newArray(slice_count); // Here, we create an array for the standard deviation values of the individual slices
median_values = newArray(slice_count); // // Here, we create an array for the standard values of the individual slices later used for background subtraction

for (i = 0; i < slice_count; i++) {
	setSlice(i+1);
	getRawStatistics(nPixels, mean, min, max, std, histogram);
	std_statistics[i] = std;
	median = getValue("Median");
	median_values[i] = median;
}

std_sorted = Array.findMaxima(std_statistics, 1); // Creates an array with sorted std values (decaying order)
focal_plane = std_sorted[0]; // Provides the slice number with the highest standard deviation

//Here, we determine the background that we want to subtract from our average bead stack
Array.getStatistics(median_values, b, b, background);

setSlice(focal_plane);
run("Enhance Contrast...", "saturated = 0.1");

// Here, we find the beads and fit them with a gaussian function to obtain the exact centroid
run("Find Maxima...", "prominence=["+peak_prominence+"] exclude output=[Point Selection]");
run("GaussFit OnSpot", "shape=Circle fitmode=[Levenberg Marquard] rectangle=["+scaled_width/2+"] pixel=1 max=500 cpcf=1 base=["+background+"]");

// Creates an overlay of focus plane and detected PSFs for quality check
selectImage(scaled_stack);
run("Flatten");
detected_peaks = getImageID();
saveAs("PNG", path + "detected_peaks.png");
run("Close");

// Now, we select and extract the PSFs of the selected fiducials
selectImage(scaled_stack);

for (i = 0; i < nResults(); i++) {
    X = getResult("X", i);
    Y = getResult("Y", i);
    run("Specify...", "width=["+scaled_width+"] height=["+scaled_width+"] x=["+X+"] y=["+Y+"] centered");
    run("Add to Manager");
}

nPSFs = roiManager("Count");

for (b=0; b<nPSFs; b++) {
	selectImage(scaled_stack);
	roiManager("Select", b);
	PSF_count = b+1;
	title = "PSF_" + PSF_count + ".tif";
	run("Duplicate...", "title=["+title+"] duplicate");
	PSF = getImageID();
	saveAs("Tiff", path2 + title); // saves the selected PSF in the created sub-directory
	selectImage(PSF);
	run("Close");
}

// Now, we import the individual PSFs as an image sequence
File.openSequence(path2);
rename("PSF_stack");

// Here, we convert the image sequence back into a hyperstack
run("Stack to Hyperstack...", "order=xyczt(default) channels=1 slices=["+slices+"] frames=["+nPSFs+"] display=Color");run("Re-order Hyperstack ...", "channels=[Channels (c)] slices=[Frames (t)] frames=[Slices (z)]");

// This section creates and saves the standard deviation image of the average PSF
if (save_PSF_std == true) {
	run("Z Project...", "projection=[Standard Deviation] all");
	PSF_std = getImageID();
	saveAs("TIFF", path + "std_PSF.tif");
	selectImage(PSF_std);
	run("Close");
	selectWindow("PSF_stack");
}

// Here we create the average PSF
run("Z Project...", "projection=[Average Intensity] all");
avg_PSF = getImageID();

// Now we have to re-order the hyperstack
run("Re-order Hyperstack ...", "channels=[Channels (c)] slices=[Frames (t)] frames=[Slices (z)]");
avg_PSF = getImageID();

// This section subtracts the average median value of all planes from the entire stack
if (subtract_background == true) {
	selectImage(avg_PSF);
	run("Subtract...", "value=["+background+"]");
}

// Finally the average PSF is saved in the selected directory
saveAs("TIFF", path + "avg_PSF.tif");


// Here, we close all the images that we created and reset the ROI manager

selectWindow("PSF_stack");
run("Close");

selectImage(scaled_stack);
run("Close");


selectImage(stack);
run("Close");

selectImage(avg_PSF);
run("Close");


if(isOpen("Results")) {
	selectWindow("Results");
	run("Close");
}

roiManager("Reset");
```
