## Supplementary material for "Transertion and cell geometry organize the *Escherichia coli* nucleoid during rapid growth": Macro_M2 - Smoothing of cell outlines in NR PAINT images.ijm

```
// This macro smoothes the membrane channel of triple-color SMLM images
// Single cells can be selected and added to the ROI manager for further analysis (e.g. erosion analysis)
// Scripted in ImageJ-macro language

// This macro is part of the publication: " Transertion and cell geometry organize the Escherichia coli nucleoid during rapid growth "

// Author:	Christoph Spahn
// 		MPI for terrestrial microbiology, Marburg
// 	


membrane_channel = 2 //define the membrane channel which is intended to be smoothed

roiManager("Reset");
run("8-bit");
membrane = getImageID();
setSlice(membrane_channel);
run("Duplicate...", " ");
mask = getImageID();
run("Gaussian Blur...", "sigma=4");
run("Auto Local Threshold", "method=Sauvola radius=30 parameter_1=0 parameter_2=0 white");
run("Fill Holes");
run("Erode");
run("Erode");
run("Gaussian Blur...", "sigma=5");
setAutoThreshold("Otsu dark");
run("Make Binary");
run("Erode");
run("Erode");
run("Erode");
run("Gaussian Blur...", "sigma=5");
setAutoThreshold("Otsu dark");
run("Make Binary");
run("Erode");
run("Erode");
run("Erode");
run("Gaussian Blur...", "sigma=5");
setAutoThreshold("Otsu dark");
run("Make Binary");
run("Dilate");
run("Dilate");
run("Gaussian Blur...", "sigma=3");
setAutoThreshold("Otsu dark");
run("Make Binary");
run("Dilate");
run("Dilate");
run("Dilate");
run("Tile");
selectImage(mask);
setTool("Wand");
roiManager("Show All with Labels");

//Select cell masks desired and add them to the RoiManager 
```
