## Supplementary material for "Transertion and cell geometry organize the *Escherichia coli* nucleoid during rapid growth": Macro_M3 - Rotation, alignment and cell straightening - CLSM.ijm

```
// This macro rotates, alignes and straightens E. coli cells acquired using confocal microscopy
// A triple-color image is required, in which the first channel represents the membrane signal
// ROIs of single bacteria of interest have already been added to the RoiManager
// Scripted in ImageJ-macro language

// This macro is part of the publication: " Transertion and cell geometry organize the Escherichia coli nucleoid during rapid growth "

// Author: 	Christoph Spahn
// 		MPI for terrestrial microbiology, Marburg
// 	

run("Conversions...", " ");
title1 = getTitle();
title2 = substring(title1, 0, lengthOf(title1)-4);
title3 = title2 + "_RoiSet.zip";
idComposite = getImageID();
run("Set Measurements...", "area centroid bounding fit redirect=None decimal=3");
run("Set Scale...", "distance=0 known=0 pixel=1 unit=pixel");

//Define Results directory -- avoid using the directory of the raw images
path = getDirectory("Choose destination for single bacteria");

run("Clear Results");
roiManager("Deselect");
ROI_loc = path + title3;
roiManager("Save", path + title3);
initROIs = roiManager("count");

for (p=0; p<initROIs; p++) {
   number = p+1;
   title4 = title2 + "_bac_" + number + "_mask.tif";
   title5 = title2 + "_bac_" + number + "_straightened.tif";
   run("Clear Results");
   roiManager("Reset");
   roiManager("Open", path + title3);
   roiManager("Deselect");
   roiManager("select", p);
   run("Measure");
   X = getResult("X");
   Y = getResult("Y");
   Winkel = getResult("Angle"); 
   Width = getResult("Width") + 10;
   Height = getResult("Height") + 10;
   run("Clear Results");
   run("Specify...", "width=Width height=Height x=X y=Y slice=1 centered");
   selectImage(idComposite);
   run("Duplicate...", "duplicate");
   StackID = getImageID();

   // Insert scaling factors here to obtain images with 10 nm pixel size
   run("Scale...", "x=8.43 y=8.43 z=1 depth=1 interpolation=Bicubic create title=scaled_ROI");
   
   run("Rotate... ", "angle=Winkel grid=1 interpolation=Bicubic enlarge stack"); 
   scaled = getImageID();
   newImage("stack", "16-bit Black", 3000, 3000, 4);
   finalStack = getImageID();
   selectImage(scaled);
   run("Copy");
   selectImage(finalStack);
   run("Paste");
   selectImage(scaled);
   setSlice(2);
   run("Copy");
   selectImage(finalStack);
   setSlice(2);
   run("Paste");
   selectImage(scaled);
   setSlice(3);
   run("Copy");
   close();
   selectImage(finalStack);
   setSlice(3);
   run("Paste");
   
   
   // This block determines cell outlines from interpolated membrane image for cell alignment on the centroid
   // Outlines are smoothed using an interative procedure of gaussian filtering, image erosion and dilation
   selectImage(StackID);
   // Insert scaling factors here to obtain images with 10 nm pixel size
   run("Scale...", "x=8.43 y=8.43 z=1.0 depth=1 interpolation=Bicubic create title=blubb");
   outline = getImageID();
   run("Rotate... ", "angle=Winkel grid=1 interpolation=Bicubic enlarge stack");
   outline2 = getImageID(); 
   selectImage(outline2);
   setSlice(1);
   run("Copy");
   run("Close");
   selectImage(finalStack);
   setSlice(4);
   run("Paste");
   run("Specify...", "width=2000 height=300 x=1500 y=1500 centered slice=1");
   run("Crop");
   setSlice(4);
   run("Duplicate...", "title=asdgda");
   blubb = getImageID();
   run("Gaussian Blur...", "sigma=1");
   setAutoThreshold("Otsu dark");
   run("Make Binary");
   run("Fill Holes");
   run("Erode");
   run("Erode");
   run("Gaussian Blur...", "sigma=5");
   setAutoThreshold("Otsu dark");
   run("Make Binary");
   run("Erode");
   run("Erode");
   run("Erode");
   run("Gaussian Blur...", "sigma=5");
   setAutoThreshold("Otsu dark");
   run("Make Binary");
   run("Erode");
   run("Erode");
   run("Erode");
   run("Gaussian Blur...", "sigma=5");
   setAutoThreshold("Otsu dark");
   run("Make Binary");
   run("Dilate");
   run("Dilate");
   run("Gaussian Blur...", "sigma=3");
   setAutoThreshold("Otsu dark");
   run("Make Binary");
   run("Dilate");
   run("Dilate");
   run("Dilate");
   run("Analyze Particles...", "size=10000-Infinity show=Masks display exclude clear include");
   mask = getImageID();
   run("Invert LUT");
   
   //This block splits the cell into ~ 300 nm thicks segments 
   
   steps = floor(getResult("Width")/30);
   start = getResult("BX");
   setForegroundColor(0,0,0);
   run("Line Width...", "line=1");
   for (ii=1; ii<=steps; ii++) {
   drawLine(start + ii*30,0,start + ii*30,300);
		}

// This block straightens the cell according to the centroids of each segment

roiManager("Reset");
run("Analyze Particles...", "size=1000-Infinity show=Nothing display exclude clear include add");
//n = nResults; 
run("Clear Results");
b = roiManager("Count");
for (f=0; f<b; f++) {
	    roiManager("select", f);
	    roiManager("Measure");
	    xcoord = getResult("BX")/10000;
	    roiManager("select", f)
        roiManager("Rename", xcoord);
        run("Clear Results");
		}
roiManager("sort");
roiManager("Deselect");
roiManager("Measure");
n = nResults;
xpoints = newArray(n);
ypoints = newArray(n); 
  for (i=0; i<n; i++) {
  	xpoints[i] = getResult("X", i);
     	ypoints[i] = getResult("Y", i);
  	}
WIDTH = getWidth();
HEIGHT = getResult("Y", n-1);
lastx = newArray(1);
lastx[0] = WIDTH;
lasty = newArray(1);
lasty[0] = HEIGHT;
firstx = newArray(1);
firstx[0] = 0;
firsty = newArray(1);
firsty[0] = getResult("Y", 0);
bla = Array.concat(firstx,xpoints,lastx);
bli = Array.concat(firsty,ypoints,lasty);

selectImage(mask);
setTool("polyline");
makeSelection("line", bla, bli);
run("Fit Spline");
roiManager("Reset");
run("Add to Manager");
selectImage(mask);
roiManager("Select", 0);
run("Flatten");
flat = getImageID();
saveAs("TIFF", path + title4);
selectImage(flat);
close();
selectImage(mask);
close();
selectImage(finalStack);

//This block performs cell straightening, line width is set to 3 �m (300 px), but can be adjusted

setSlice(1);
run("Duplicate...", " ");
membrane = getImageID();
roiManager("Select", 0);
run("Straighten...", "title=membrane line=300");
membrane_straight = getImageID();
rename("membrane");
run("Green");
selectImage(membrane);
close();
selectImage(finalStack);
setSlice(2);
run("Duplicate...", " ");
MreB = getImageID();
roiManager("Select", 0);
run("Straighten...", "title=MreB line=300");
MreB_straight = getImageID();
rename("MreB");
run("Red Hot");
selectImage(MreB);
close();
selectImage(finalStack);
setSlice(3);
run("Duplicate...", " ");
DNA = getImageID();
selectImage(finalStack);
close();
selectImage(DNA);
roiManager("Select", 0);
run("Straighten...", "title=DNA line=300");
DNA_straight = getImageID();
run("Cyan Hot");
selectImage(DNA);
close();
selectImage(blubb);
close(); 
selectImage(DNA_straight);
rename("DNA");

//This block creates the composite and determines the cell length used in the filename to sort cells by length

run("Merge Channels...", "c1=membrane c2=MreB c3=DNA create");
selectWindow("Composite");
straight_3C = getImageID();
run("Duplicate...", "title=asdgda");
Outlines = getImageID();
setAutoThreshold("Otsu dark");
run("Make Binary");
run("Fill Holes");
run("Analyze Particles...", "size=20000-Infinity circularity=0.00-1.00 show=Nothing display clear include record");
XC = 1000 - getResult("X");
YC = 150 - getResult("Y");
width = getResult("Width")/100;
run("Close");
selectImage(straight_3C);
run("Translate...", "x=XC y=YC interpolation=None stack");
run("Clear Results");
bacsize = d2s(width, 2); 
title6 = bacsize + "_" + title5;
run("16-bit");
saveAs("TIFF", path + title6);
selectImage(straight_3C);
close(); 
selectImage(StackID);
close();
roiManager("Reset");
}
run("Clear Results");
selectWindow("Results");
run("Close");
```
