## Supplementary material for "Transertion and cell geometry organize the *Escherichia coli* nucleoid during rapid growth": Macro_M4 - Cell normalization - CLSM.ijm

```
// This macro normalizes 3Ch confocal images of single cells to a fixed length and width
// .tiff files of single bacteria need to be placed in the same folder
// Scripted in ImageJ-macro language

// This macro is part of the publication: " Transertion and cell geometry organize the Escherichia coli nucleoid during rapid growth "

// Author: 	Christoph Spahn
// 		MPI for terrestrial microbiology, Marburg
// 	

dir1 = getDirectory("Choose Source Directory ");
dir2 = getDirectory("Choose Destination Directory "); //other than source directory
list = getFileList(dir1);
run("Set Measurements...", "area centroid bounding fit redirect=None decimal=3");
setBatchMode(true);
for (i=0; i<list.length; i++) {
	open(dir1+list[i]);
	id = getImageID();
	title = getTitle();
	title2 = substring(title, 0, lengthOf(title)-4);
	title3 = title2 + "_normalized_l_w";
	setSlice(1);
	run("Duplicate...", "title=cell_width");
	dupl = getImageID();
	duplicate_width = getImageID();
	run("Gaussian Blur...", "sigma=2");
	setAutoThreshold("Otsu dark");
	run("Make Binary");
	run("Analyze Particles...", "size=20000-500000 circularity=0.1-1.00 display exclude clear include");
	BX = getResult("BX");
	BY = getResult("BY");
	Width = getResult("Width");
	Height = getResult("Height");	
	selectImage(dupl);
	run("Close");
	selectImage(id);
	makeRectangle(BX, BY, Width, Height);
	run("Crop");
	crop = getImageID();
	run("Scale...", "x=- y=- z=1.0 width=500 height=150 depth=3 interpolation=Bicubic average process create title=bla");
	saveAs("TIFF", dir2+ title3);
	selectImage(crop);
	close();
}
setBatchMode(false);
```
