## Supplementary material for "Transertion and cell geometry organize the *Escherichia coli* nucleoid during rapid growth": Macro_M5 - Determination of the relative nucleoid length expansion (RNLE) - CLSM.ijm

```
// This macro determines the cell and nucleoid width in single cells
// All single-cell .tiff files of straightened cells need to be in one directory 
// Results are summarized in a table and saved in the source directory
// Scripted in ImageJ-macro language

// This macro is part of the publication: " Transertion and cell geometry organize the Escherichia coli nucleoid during rapid growth "

// Author: 	Christoph Spahn
// 		MPI for terrestrial microbiology, Marburg
// 	

dir1 = getDirectory("Choose Source Directory ");
list = getFileList(dir1);
title3 = "Lengths_and_nucleoid_expansion";
titleTable = "["+title3+"]";
f=titleTable;

// This block creates the results table
run("New... ", "name="+titleTable+" type=Table"); 
print(f,"\\Headings:cell\tLenght\tnucleoid_width"); 

// This block extracts the cell length from the title and determines the nucleoid length
run("Set Measurements...", "area centroid bounding fit shape integrated redirect=None decimal=3");
for (i=0; i<list.length; i++) {
   showProgress(i+1, list.length);
   open(dir1+list[i]);
   title = getTitle();
   title2 = substring(title, 0, lengthOf(title)-4);
   id = getImageID();
   setSlice(1);
   run("Duplicate...", " ");
   Outlines = getImageID();
   bacsize = substring(title, 0, 4);
   run("Gaussian Blur...", "sigma=2");
   setAutoThreshold("Otsu dark");
   run("Make Binary");
   run("Analyze Particles...", "size=20000-Infinity circularity=0.00-1.00 show=Nothing exclude clear include add");
   run("Set Measurements...", "area bounding redirect=None decimal=3");
   selectImage(Outlines);
   close();
   selectImage(id);
   setSlice(3);
   run("Duplicate...", " ");
   DNA_full = getImageID();
   setAutoThreshold("Moments dark");   
   run("Make Binary");   
   roiManager("select", 0);
   run("Duplicate...", " ");
   DNA = getImageID();
   selectImage(DNA_full);
   close();
   selectImage(DNA);
   roiManager("Reset");
   run("Analyze Particles...", "size=200-Infinity circularity=0.10-1.00 show=Nothing clear include add");
   run("Clear Results");
   getDimensions(width, height, X, X, X);
   newImage("nucleoids", "8-bit black", width, height, 1);
   nucleoid_width = getImageID();
   selectImage(DNA);
   close();
   selectImage(nucleoid_width);
   roiManager("Deselect");
   setForegroundColor(255, 255, 255);
   roiManager("Fill");
   roiManager("Reset");
   run("Scale...", "x=- y=- width=width height=1 interpolation=None average create title=bla");
   blubb = getImageID();
   setThreshold(1, 255);
   run("Convert to Mask");
   run("Divide...", "value=255");
   run("Set Measurements...", "area bounding integrated redirect=None decimal=3");
   run("Measure");
   nucleoid = getResult("RawIntDen")/100;
   selectImage(nucleoid_width);
   close();
   selectImage(blubb);
   close();
   selectImage(id);
   close();
   cell = i+1;
   print(f,cell+"\t"+bacsize+"\t"+nucleoid);
   run("Clear Results");
}
selectWindow(title3);
title4 = title3 + ".txt";
saveAs("Results", dir1+ title4);
selectWindow(title3);
run("Close");  		
roiManager("reset");
```
