## Supplementary material for "Transertion and cell geometry organize the *Escherichia coli* nucleoid during rapid growth": Macro_M6 - Determination of the relative MreB distribution - CLSM.ijm

```
// This macro determines the MreB intensity in confocal average images
// Scripted in ImageJ-macro language

// This macro is part of the publication: " Transertion and cell geometry organize the Escherichia coli nucleoid during rapid growth "

// Author: 	Christoph Spahn
// 		MPI for terrestrial microbiology, Marburg
// 	


dir1 = getDirectory("Open results directory"); //destination for ROIs and results tables
roiManager("Reset");
run("Clear Results");
run("Input/Output...", "jpeg=80 gif=-1 file=.txt save_column");
raw = getImageID();
title = getTitle();
title2 = substring(title, 0, lengthOf(title)-4) + "_RoiSet.zip";
title3 = substring(title, 0, lengthOf(title)-4) + "_mask.tif";
title4 = substring(title, 0, lengthOf(title)-4) + "_results.txt";
setForegroundColor(0, 0, 0);
run("Input/Output...", "jpeg=80 gif=-1 file=.txt save_column");
run("Line Width...", "line=1");
run("Set Measurements...", "area bounding redirect=None decimal=3");
setSlice(1);
run("Duplicate...", " ");
ROIs = getImageID();
setAutoThreshold("Otsu dark");
run("Make Binary");
run("Duplicate...", " ");
cytosol = getImageID();
run("Invert");
run("Analyze Particles...", "size=1000-Infinity display exclude clear add");
f = roiManager("Count");
if (f>=2) {
	run("Clear Results");
	roiManager("Select", newArray(0,1));
	roiManager("Combine");
	roiManager("Delete");
	roiManager("Add");
	roiManager("Measure");
}
roiManager("Reset");
selectImage(cytosol);
run("Close");
selectImage(ROIs);
//run("Measure");
BX = getResult("BX");
width = getResult("Width");
BX2 = BX + width;
drawLine(BX, 0, BX, 150);
drawLine(BX2, 0, BX2, 150);
saveAs("TIFF", dir1 + title3);

// Pole and cylinder regions need to be added to the ROI manager
// Add both poles as one ROI using the shift key

roiManager("Reset");
run("Clear Results");
run("Set Measurements...", "area integrated redirect=None decimal=3");
waitForUser("ROI selection", "Add ROIs of pole regions and cylindrical sections");
roiManager("Select", newArray(0,1));
roiManager("XOR");
roiManager("Add");

roiManager("Select", 0);
roiManager("Rename", "poles");
roiManager("Select", 1);
roiManager("Rename", "cylinder");
roiManager("Select", 2);
roiManager("Rename", "all");
roiManager("Save", dir1 + title2);
selectImage(ROIs);
run("Close");

selectImage(raw);
run("Slice Remover", "first=1 last=3 increment=2");
MreB = getImageID();
roiManager("Deselect");
roiManager("Measure");
selectWindow("Results");
saveAs("Results", dir1 + title4);
selectWindow("Results");
run("Close");
roiManager("Reset");
selectImage(MreB);
run("Close");

// The text file need to be saved separately
```
