## Supplementary material for "Transertion and cell geometry organize the *Escherichia coli* nucleoid during rapid growth": Macro_M7 - Determination of radial intensity distributions using erosion analysis.ijm

```
// This macro erodes the outlines of single bacteria in order to measure the membrane- and DNA-intensity of the resulting segments;
// Required is a 3Ch image, that contains both a PAINT channel of the DNA and membrane and a diffraction limited widefield channel;
// The desired ROIs need to be opened, same for the source image
// Scripted in ImageJ-macro language

// This macro is part of the publication: " Transertion and cell geometry organize the Escherichia coli nucleoid during rapid growth "

// Author: 	Christoph Spahn
// 		MPI for terrestrial microbiology, Marburg
// 	


// Define image channels
widefield_channel = 1;
membrane_channel = 2;
DNA_channel = 3;


idComposite = getImageID();
path = getDirectory("Destination single bacteria");
run("Clear Results");
title1 = getTitle();
title2 = substring(title1, 0, 2); //if number of measurements > 10, change to 0, 3! 
title3 = title2 + "_RoiSet.zip";
roiManager("Save", path + title3);
initROIs = roiManager("count");
setSlice(3);
run("Add Slice", "add=channel");
setSlice(4);
resetMinAndMax();
roiManager("Deselect");
setForegroundColor(255,255,255);
roiManager("Fill");
four_channels = getImageID();

for (p=0; p<initROIs; p++) {
	run("Set Measurements...", "area centroid bounding fit integrated redirect=None decimal=3");
	selectImage(four_channels);
	number = p+1;
	title4 = title2 + "_bac_" + number + "_mask.tif";
	title5 = title2 + "_bac_" + number + "_erosion_ROIs.zip";
	title6 = title2 + "_bac_" + number + "_3Ch_single_bac.tif";
	title7 = title2 + "_bac_" + number + "_DNA_erosion.txt";
	title8 = title2 + "_bac_" + number + "_membrane_erosion.txt";
	title9 = title2 + "_bac_" + number + "_single_bac_erosion_ROIs.png";
	title10 = title2 + "_bac_" + number + "_rel_area_DNA_membrane_int_statistics.txt";
	run("Clear Results");
	
	//this block duplicates the single bacterias and erodes the membrane mask for radial segmentation

	roiManager("Deselect")
	roiManager("select", p)
	run("Measure");
	X = getResult("BX")-10;
	Y = getResult("BY")-10;
	Width = getResult("Width") + 20;
	Height = getResult("Height") + 20;
	run("Clear Results");
	run("Specify...", "width=Width height=Height x=X y=Y slice=1"); 
	run("Duplicate...", "title=[bla] duplicate channels=1-4");
	single_bac = getImageID();
	setSlice(4);
	run("Duplicate...", " ");
	mask = getImageID();
	roiManager("Reset");	

	run("Analyze Particles...", "size=500-Infinity circularity=0.10-1.00 exclude include add");
	roiManager("Select", 0);
	setBackgroundColor(0,0,0);
	run("Clear Outside");
	run("Select None");
	n = 30;
	for (i=0; i<n; i++) {
		nROI_1 = roiManager("Count");
		run("Erode");
		run("Erode");
		run("Erode");
		run("Erode"); //results in 80nm erosion steps
		//run("Erode"); //results in 100 nm erosion steps
		run("Analyze Particles...", "size=500-Infinity circularity=0.00-1.00 exclude include add");
		nROI_2 = roiManager("Count");
		if (nROI_2-nROI_1 > 1) {
			add_ROI = nROI_2 - nROI_1;
			combined_ROIs = newArray(add_ROI);
			for (h=0; h<add_ROI; h++) {
				combined_ROIs[h] = nROI_1+h;	
				roiManager("Select", combined_ROIs);
			}
			roiManager("Combine");
			roiManager("Add");
			roiManager("Select", combined_ROIs);
			roiManager("Delete");
			run("Select None");
		}
		}

	roiManager("Save", path + title5);
	selectImage(mask);
	close();
	selectImage(single_bac);
	setSlice(membrane_channel);
	run("Clear Results");
	roiManager("Deselect");
	run("Set Measurements...", "area fit integrated redirect=None decimal=3");
	roiManager("Measure");
	saveAs("Results", path+ title8);

	//this block creates a table that saves the relative areas and relative intensities
	results = nResults;
	tabtitle = "statistics";
	titleTable = "["+tabtitle+"]";
	m=titleTable;
	run("New... ", "name="+m+" type=Table"); 
	print(m,"\\Headings:Area\trelative_area\trelative_membrane_intensity\trelative_DNA_intensity");
	results = nResults;
	disc_area = newArray(results+1);
	rel_area = newArray(results+1);
	rel_int_DNA = newArray(results+1);
	rel_int_membrane = newArray(results+1);

	//this block determines the relative area and relative membrane intensity of the eroded disks
	for (aa=0; aa<results; aa++) {
	disc_area[aa] = getResult("Area", aa);
	rel_area[aa] = disc_area[aa]/disc_area[0];
	//print(disc_area[aa], rel_area[aa]);
	}
	for (bb=0; bb<results; bb++) {
		if (bb+1<nResults) {
		rel_int_membrane[bb] = (getResult("RawIntDen", bb)-getResult("RawIntDen", bb+1))/getResult("RawIntDen", 0);
		} else {
			rel_int_membrane[bb] = getResult("RawIntDen", bb)/getResult("RawIntDen", 0);
		}
		//print(rel_int_membrane[bb]);
	}
	
	selectImage(single_bac);
	setSlice(DNA_channel);
	roiManager("Deselect");
	run("Clear Results");
	roiManager("Measure");
	saveAs("Results", path+ title7);

	//this block determines the relative DNA intensity of the eroded disks
	for (cc=0; cc<results; cc++) {
		if (cc+1<nResults) {
		rel_int_DNA[cc] = (getResult("RawIntDen", cc)-getResult("RawIntDen", cc+1))/getResult("RawIntDen", 0);
		} else {
			rel_int_DNA[cc] = getResult("RawIntDen", cc)/getResult("RawIntDen", 0);
		}
		//print(rel_int_DNA[cc]);
	print(m,disc_area[cc]+"\t"+rel_area[cc]+"\t"+rel_int_membrane[cc]+"\t"+rel_int_DNA[cc]);
	}

	//this block fills the table and saves it in the results_directory
	//print(m,disc_area[aa]+"\t"+rel_area[aa]+"\t"+rel_int_membrane[bb]+"\t"+rel_int_DNA[cc]);
	selectWindow(tabtitle);
	saveAs("Results", path+ title10);
	selectWindow(tabtitle);
	run("Close");
	
	selectImage(single_bac);
	roiManager("Deselect");
	run("Duplicate...", "title=[bli] duplicate channels=1-3");
	single_bac_3Ch = getImageID();
	saveAs("TIFF", path + title6);
	setForegroundColor(255, 255, 0);
	selectImage(single_bac_3Ch);
	roiManager("Set Color", "yellow");
	roiManager("Set Line Width", 1);
	roiManager("Deselect");
	roiManager("Draw");
	saveAs("PNG", path + title9); 
	selectImage(single_bac_3Ch);
	close();
	selectImage(single_bac);
	close();
	roiManager("Reset");
	roiManager("Open", path + title3);
}
selectImage(four_channels);
run("Close");
roiManager("Reset");
run("Clear Results");
selectWindow("Results");
run("Close");
```
