## Supplementary material for "Transertion and cell geometry organize the *Escherichia coli* nucleoid during rapid growth": Macro_M8 - Plot profiles along perimenter.ijm

```
// This macro is used to plot intensity profiles along the cell perimeter of super-resolved cells
// Requirements are multichannel images with at least 2 channels (DNA and MreB)

// This macro is part of the publication: " Transertion and cell geometry organize the Escherichia coli nucleoid during rapid growth "

// Author: 	Christoph Spahn
// 		MPI for terrestrial microbiology, Marburg
// 	

// This macro was written in Fiji version 1.53q


// Variables
MreB_channel = 1;
DNA_channel = 3;
dilation_radius = -20; // This parameter defines how many pixels ROIs are enlarged or shrinked; Negative numbers = shrinking
line_width = 30; // This parameter defines the line thickness for the plot
pixel_size = 0.01 // Define the pixel size in µm

run("Input/Output...", "jpeg=100 gif=-1 file=.txt use_file copy_row save_column save_row");

dir = getDirectory("Choose output directory");
img = getImageID();
title = getTitle();
title2 = title.replace(".tif", "_updated_ROIs.zip");
run("Set Scale...", "distance=1 known=["+pixel_size+"] unit=µm");


// Macro

// Count the number of ROIs for this measurement
nRois = roiManager("Count");
roi_count = 0


// This section converts the ROIs to a polyline ROI
roi_selection = newArray(nRois);

for (i = 0; i < nRois; i++) {
	roi_selection[i] = i;
	roiManager("Select", i);
	run("Enlarge...", "enlarge=["+dilation_radius+"] pixel");
	//run("Fit Spline");
	roiManager("Update");
	run("Area to Line");
	roiManager("Update");
	getSelectionCoordinates(xpoints, ypoints);
	firstx = xpoints[0];
	firsty = ypoints[0];
	xpoints2 = Array.concat(xpoints,firstx);
	ypoints2 = Array.concat(ypoints,firsty);
	makeSelection("polyline", xpoints2, ypoints2);
	roiManager("add");
}

roiManager("Select", roi_selection);
roiManager("Delete");

run("Line Width...", "line=["+line_width+"]");

setBatchMode(true);
for (i = 0; i < nRois; i++) {
	// This part creates a new table, in which the profile values of DNA and MreB plots are saved
	extension = "_ROI_" + i;
	title3= title.replace(".tif", extension);
	titleTable = "["+title3+"]";
	f=titleTable;
	
	run("New... ", "name="+titleTable+" type=Table"); 
	print(f,"\\Headings:distance\tMreB_signal\tDNA_signal"); 

	roiManager("Select", i);
	
	// This block gets the intensity values for the MreB and DNA channel
	run("Clear Results");
	selectImage(img);
	setSlice(MreB_channel);
	run("Plot Profile");
	Plot.getValues(xpoints, ypoints);
	wait(500);
	run("Close");
	selectImage(img);
	setSlice(DNA_channel);
	run("Plot Profile");
	Plot.getValues(xpoints2, ypoints2);
	wait(500);
	run("Close");
	
	for (b = 0; b < xpoints.length; b++) {
		print(f,xpoints[b]+"\t"+ypoints[b]+"\t"+ypoints2[b]);
	}

	// Now we save the table in the selected directory and close it afterwards
	selectWindow(title3);
	title4 = title3 + ".txt";
	saveAs("Results", dir+ title4);
	selectWindow(title3);
	run("Close");  
	
	run("Clear Results");
	
	roi_count = roi_count+1;
}
setBatchMode(false);

roiManager("Deselect");
roiManager("Save", dir + title2);

roiManager("Reset");

selectImage(img);
run("Close");

print("Profiles of " + roi_count + " cells were measured with a shrinking radius of " + dilation_radius + " pixels and a profile width of " + line_width + " pixels");
```
