## Supplementary material for "Transertion and cell geometry organize the *Escherichia coli* nucleoid during rapid growth": Macro_M9 - Simulate images for circular cross-correlation.ijm

```
// This macro is used to simulate 3-color images for circular cross-correlation
// It positions point-like signals equidistantly along a circle with a defined rotational shift
// Scripted in ImageJ-macro language

// This macro is part of the publication: " Transertion and cell geometry organize the Escherichia coli nucleoid during rapid growth "

// Author:	Christoph Spahn
// 		MPI for terrestrial microbiology, Marburg
// 	

// This macro was written in Fiji version 1.53q


save_stuff = true; // set true if you want to automatically save the simulated image and results

roiManager("Reset");

if (isOpen("Log")) {
	selectWindow("Log");
	run("Close");
}

// These are the image parameters

img_width = 500;
img_height = 500;

// These parameters define the circle

r = 200; // radius in pixels
h = 250; // circle center x
k = 250; // circle center k

x_circ = h-r;
y_circ = k-r;
circ_width = 2*r;
circ_height = 2*r;

blur = 4; // Sigma of the Gaussian Blur of the final image in px

// Here we create the image

newImage("Untitled", "32-bit black", img_width, img_height, 3);

stack = getImageID();

// define the number of points and the increment in first fluorescent channel and the rotational shift of the second channel

Ch1_npoints = 12;
Ch2_npoints = 12;
Ch1_shift = 80; // rotational shift in degree for Ch1
Ch2_shift = 80; // rotational shift in degree for Ch2

Ch1_rad_shift = Ch1_shift * 0.0174533;
Ch2_rad_shift = Ch2_shift * 0.0174533;

Ch1_rad_diff = (2*PI)/Ch1_npoints;
Ch2_rad_diff = (2*PI)/Ch2_npoints;

Ch1_disp = 10; // Displacement of points in Ch1 towards center in px
Ch2_disp = 30; // Displacement of points in Ch2 towards center in px


run("Line Width...", "line=1");

// Here we draw the oval
makeOval(x_circ, y_circ, circ_width, circ_height);
roiManager("Add");
roiManager("Select", 0);
roiManager("Rename", "Oval");


// Here we draw the the circle and points

selectImage(stack);
setSlice(1);

setForegroundColor(255, 255, 255);
roiManager("Select", 0);
roiManager("Draw");

// Now we create the point selection for channel 1

Ch1_x = newArray(Ch1_npoints);
Ch1_y = newArray(Ch1_npoints);

for (i = 0; i < Ch1_npoints; i++) {
	t = Ch1_rad_diff*i + Ch1_rad_shift;
	Ch1_x[i] = points_on_circle_x(r-Ch1_disp, h, k, t);
	Ch1_y[i] = points_on_circle_y(r-Ch1_disp, h, k, t);
}

selectImage(stack);
setSlice(2);
makeSelection("multipoint", Ch1_x, Ch1_y);
roiManager("Add");
roiManager("Select", 1);
roiManager("Rename", "Ch1_points");
roiManager("Draw");


// Now we create the point selection for channel 1

Ch2_x = newArray(Ch2_npoints);
Ch2_y = newArray(Ch2_npoints);

for (i = 0; i < Ch2_npoints; i++) {
	t = Ch2_rad_diff*i+ Ch2_rad_shift;
	Ch2_x[i] = points_on_circle_x(r-Ch2_disp, h, k, t);
	Ch2_y[i] = points_on_circle_y(r-Ch2_disp, h, k, t);
}

selectImage(stack);
setSlice(3);
makeSelection("multipoint", Ch2_x, Ch2_y);
roiManager("Add");
roiManager("Select", 2);
roiManager("Rename", "Ch2_points");
roiManager("Draw");

run("Re-order Hyperstack ...", "channels=[Slices (z)] slices=[Channels (c)] frames=[Frames (t)]");

stack = getImageID();

setSlice(1);
run("Gaussian Blur...", "sigma=["+blur+"]");
run("Red");
run("Enhance Contrast...", "saturated=0.1");

setSlice(2);
run("Gaussian Blur...", "sigma=["+blur+"]");
run("Cyan Hot");
run("Enhance Contrast...", "saturated=0.1");

setSlice(3);
run("Gaussian Blur...", "sigma=["+blur+"]");
run("Yellow Hot");
run("Enhance Contrast...", "saturated=0.1");

run("Make Composite");


function points_on_circle_x(r, h, k, t){
	x_pos = r*cos(t) + h;
	return x_pos;
}

function points_on_circle_y(r, h, k, t){
	y_pos = r*sin(t) + k;
	return y_pos;
}

print("Image size was " + img_height + "x " + img_width + " pixels.");
print("Circle radius was " + r + " pixels.");
print("For Channel 1, " + Ch1_npoints + " points were positioned in " + 360/Ch1_npoints + " degree distance and a displacement of " + Ch1_disp + " pixels from the circle, and a shift of " + Ch1_shift + " degree.");
print("For Channel 2, " + Ch2_npoints + " points were positioned in " + 360/Ch2_npoints + " degree distance, a displacement of " + Ch2_disp + " pixels from the circle, and a shift of " + Ch2_shift + " degree.");


if (save_stuff == true) {
	
	dir = getDirectory("Select folder to save images");
	
	selectImage(stack);
	saveAs("TIFF", dir + "Stack.tif");
	run("Close");
	roiManager("Deselect");
	roiManager("Save", dir + "Simulated_ROIs.zip");
	roiManager("Select", 1);
	roiManager("Delete");
	roiManager("Select", 1);
	roiManager("Delete");
	roiManager("Select", 0);
	roiManager("Save", dir + "Circle.zip");
	roiManager("Reset");
	
	selectWindow("Log");
	saveAs("Text", dir + "Simulation_parameters.txt");
	
	if (isOpen("Results")) {
		selectWindow("Results");
		run("Close");
	}
	
	if (isOpen("Log")) {
	selectWindow("Log");
	run("Close");
	}
}
```
